# Pervasive fitness trade-offs revealed by rapid adaptation to shifting population densities in large experimental populations of *Drosophila melanogaster*

**DOI:** 10.1101/2024.10.28.620721

**Authors:** M.C. Bitter, S. Greenblum, S. Rajpurohit, A.O. Bergland, J.A. Hemker, Egor Lappo, N.J. Betancourt, S. Tilk, S. Berardi, H. Oken, P. Schmidt, D.A. Petrov

## Abstract

Trade-offs are an inherent feature of organismal biology that are expected play a fundamental role in the evolution of natural populations. Efforts to quantify trade-offs are largely confined to phenotypic measurements and the identification of negative genetic-correlations among fitness-relevant traits. Here, we use time-series genomic data collected during experimental evolution in large, genetically diverse populations of *Drosophila melanogaster* to directly measure the manifestation of trade-offs in response to fluctuating selection on ecological timescales. Specifically, we first conducted a lab-based selection experiment to quantify a genome-wide signal of antagonistic pleiotropy elicited in response to shifting population densities and associated with reproduction and stress tolerance selection. In doing so, we identified a putative role of two cosmopolitan inversions in these trade-offs. We then conducted an independent experiment to show that a simple manipulation of increasing population density under controlled lab-based conditions identified loci that are relevant to selection during population expansion and collapse in a complex, semi-natural setting. In concert, our results reveal how adaptation in complex, natural environments can be coarse-grained in such a manner to drive repeatable and predictable patterns of genomic variation, and further add credence to models positing a role of generic fitness trade-offs in the maintenance of variation in natural populations.

## Introduction

A fundamental tenet of evolutionary theory is that trait adaptation is restricted by trade-offs: the cost to individual fitness when an advantageous change in one trait occurs at the detriment to another (1,2). Trade-offs can be observed across closely related species, such as those between beak and body size among Darwin’s finches (3,4). They can also arise as a consequence of microevolutionary processes among populations of the same species; for instance, the trade-offs to survival associated with distinct reproductive strategies among populations of salmon (5). Trade-offs oftentimes emerge, and are most readily studied, in the context of life-history traits, and form the basis of the theory of life history evolution (2). Ultimately, a key theoretical implication of trade-offs is that they maintain variation by constraining the simultaneous optimization of traits associated with fitness under different environmental conditions (2,6–8).

At the molecular level, trade-offs likely imply antagonistic pleiotropy, a phenomenon in which an allele that enhances one fitness-related trait may simultaneously impede another in a particular environment. Then, when the environment changes and selection favors the previously impeded trait, the previously favored allele becomes disadvantageous (7–9). Thus, a widely used method for inferring the presence of antagonistic pleiotropy and trade-offs is to compute genetic (co) variances using phenotypic data (10–12). Still another, putatively more direct, signature of trade-offs are fluctuations in population allele frequencies that occur in concert over time with selective pressures. Indeed, several recent studies observed genome-wide fluctuations in allele frequencies through time, and across several distinct systems, suggesting underlying trade-offs (13–18).

A fundamental life-history trade-off is the balance between directing energy either towards survival or reproduction whereby periods of stressful environmental conditions favor somatic maintenance at the cost of reproductive investment (19). Evidence of such a trade-off is widespread across species, from soay sheep (20), to seaweed flies (21), to humans (22). It is clearly apparent among populations of *Drosophila melanogaster* inhabiting temperate environments, a well-studied system in which increased food accessibility and warmer climates during summer spur exponential population growth, followed in winter by harsher conditions and population collapse (23–25). These cyclical boom-bust population dynamics occur over approximately 10 generations and coincide with the evolution of several classic life history traits: traits conferring increased reproduction (e.g., fecundity, faster developmental rates) are favored during spring and summer as populations expand, followed by an increase in stress tolerance traits (e.g., desiccation and starvation resistance) throughout the harsh winter and subsequent population collapse (26–28). It is likely that trade-offs underpin these contrasting patterns of adaptation, as negative genetic correlations have been quantified between seasonally evolving reproduction and stress tolerance traits (10,11,27,29–31).

Recent genome-wide sequencing of wild populations, and outbred populations evolved in semi-natural mesocosms, has revealed evidence that hundreds of independent loci exhibit signatures of selection concurrent to these patterns of rapid phenotypic evolution, many of which switch in sign in concert with the fluctuating environment and evolving traits (13–16). Strikingly, despite differences in the particular environmental conditions in which patterns of evolutionary change were assayed (e.g., different observation years and geographic locations), there is high repeatability in the genomic regions subject to fluctuating selection across space and time (13–15). One hypothesis for this predictability at the genomic level is that the loci responding to selection are not sensitive to specific environmental variables (which can be idiosyncratic across study locales and years), but rather respond more coarsely to the generic and repeatable demographic fluctuations in the system (i.e., shifts in population density). The observed parallel genomic signals of temporally varying selection in this system may then represent the manifestation of fundamental fitness trade-offs emerging in response to these shifting population densities. Systematically and rigorously testing this hypothesis, however, necessitates controlling for the complex suite of abiotic shifts that occur in outdoor environments and isolating the relative contribution of demographic selection to patterns of allele frequency change.

Here, we conducted two experimental evolution studies to systematically quantify how predictable evolutionary trajectories arise in complex, fluctuating environments and, in effect, explicitly link key components of life history evolution to patterns of fluctuating selection and maintenance of variation in *D. melanogaster*. In the first of these studies, we assayed the impact of demographic shifts, independent from any environmental selection. To do so, we monitored patterns of genomic variation in genetically diverse, replicate populations housed in a controlled, indoor environment throughout eleven discrete (non-overlapping) generations of population expansion, and a single bout of population truncation (Fig. 1; Fig S1). Our replicate populations were derived from a reconstituted, outbred population initiated from 145 inbred (DGRP) lines (Methods; Supplementary Data File 1), and evolution proceeded in four replicate cages (2 m^3^). Food availability in the replicate cages was commensurate with increasing population sizes during the expansion phase, generating a selective regime that eliminated the impact of intraspecific competition (e.g. for food or egg laying space) and favored the reproduction-associated traits expected when resources are abundant, such as fecundity and developmental rate (27). Contrarily, population truncation was induced by removing all food and water from our replicate cages and sampling the surviving individuals as the replicates collapsed. This mirrors dynamics of a population that has exceeded carrying capacity and, in effect, selects for the ability to withstand depleted resources via stress-tolerance mechanisms (32), which in our experiment is most likely to be desiccation resistance (33). We hypothesized that these contrasting selective regimes (hereafter, ‘reproduction’ and ‘stress-tolerance’ selection) would reveal trade-offs between life-history variation, whereby antagonistically pleiotropic alleles favored during sustained population expansion would become selected against during truncation.

**Figure 1.**
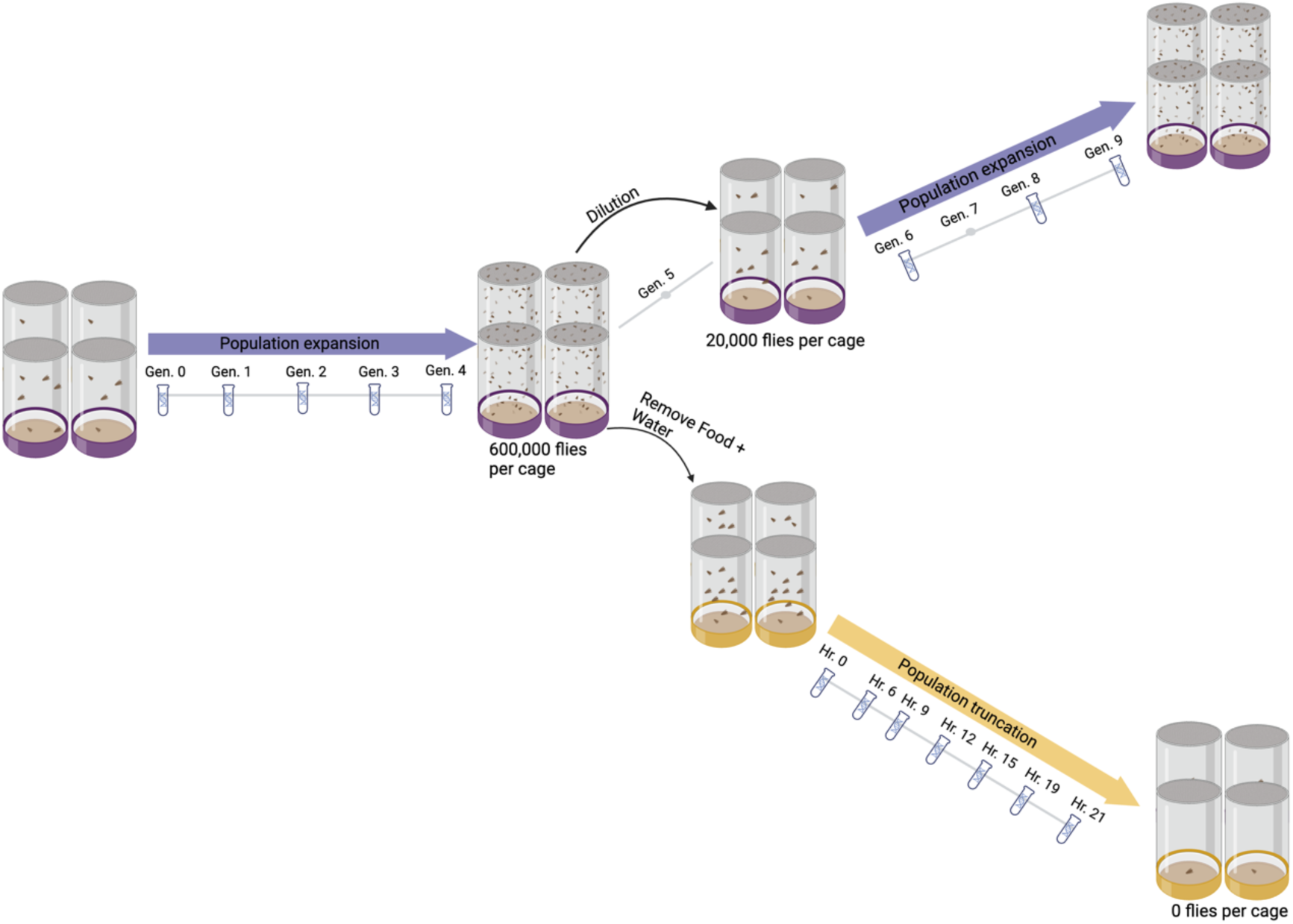
Schematic of lab-based, population expansion-truncation selection experiment. Four replicate, 2 m^3^ cages were seeded with a genetically diverse, outbred population generated via four generations of recombination (N = 145 inbred lines). Evolution proceeded in discrete generations, whereby the amount of food was doubled every generation until replicate populations reached a census size of approximately 600,000 flies per cage (generation four). At this point, eggs from the generation 5 cohort were collected and all food and water was removed from the replicate cages. The adult cohort of the generation 4 flies was then sampled at seven time points (hours 0-21) as the absence of resources collapsed the populations. The generation 5 eggs were used to re-seed the replicate cages, which underwent continued population expansion for four additional generations. DNA in vials denotes generations of expansion, and hours of truncation, during which pooled samples were collected for pooled allele frequency calculation. Schematic generated with bioRender (https://www.biorender.com/).

The second experiment leveraged an independent inbred panel to construct a genetically diverse, outbred population, which was split into a set of indoor cages (N = 10) and outdoor mesocosm (N = 12) replicates. Through this, we used the same mapping population (thereby eliminating any confounding impacts of differences in recombination and/or linkage disequilibrium) to directly test whether loci identified under selection during increasing population density in a controlled, indoor environment are relevant to selection and trade-offs during adaptation in an outdoor, naturally fluctuating environment. In concert, our results provide evidence for: (1) strong, parallel selection elicited by changes in population density (2) genome-wide tradeoffs associated with contrasting selective regimes of population expansion and truncation, (3) the role of cosmopolitan inversions underpinning fitness trade-offs in the species and (4) relevance of the loci responding to density selection in a controlled, lab-based setting to patterns of adaptation in response to natural environmental fluctuations.

## Results

### Selection during population expansion elicits genome-wide signals of strong, parallel selection

We observed parallel shifts in allele frequencies across the four replicate populations throughout the progression of population expansion, indicating adaptive responses to the shared selective pressures imposed in our experimental system. We quantified these systematic allelic shifts using several approaches. We first computed genome-wide divergence as average F_ST_ across all segregating sites pairwise among all expansion samples and used these data to carry out several tests. F_ST_ divergence from the generation 0 populations increased steadily within each cage through time, readily exceeding differentiation between biological replicates (F = 172.2; p-value < 0.01; Fig. 2A & Fig. S2). We then used pairwise F_ST_ values as a distance metric to create multi-dimensional scaling (MDS) plots, in which divergence between samples is represented as distance among the points in a 2-D plane. Coloring samples based on the replicate cage identity indicated that genome-wide allele frequencies in one replicate were perturbed, shifting its points from the remaining three replicates throughout expansion (Fig. 2B). We hypothesize this was likely a result of potential bottlenecks during replicate cage founding. However, despite this offset, samples from all cages appeared to shift across the 2-D plane in the same direction through time, suggesting that the replicate populations experienced parallel shifts in genome-wide allele frequencies (Fig 2C and Fig S3; a trend that was also detected using principal component analysis; Fig. S4).

**Figure 2.**
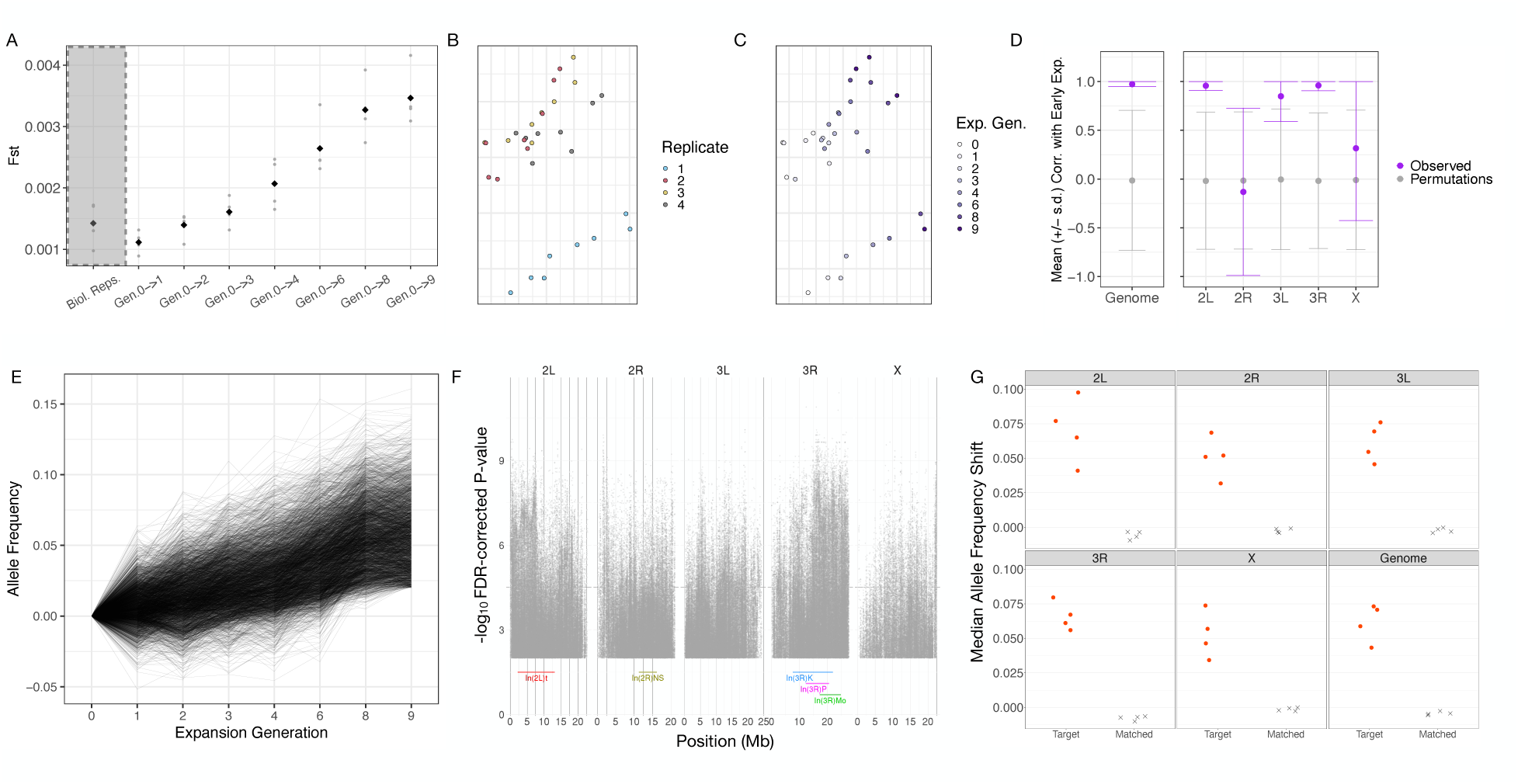
Adaptation under sustained population expansion elicits strong, genome-wide parallel responses. (A) Mean, genome-wide F_ST_ between biological replicates (‘Biological Replicates’; same replicate/collection time point, different pooled sample/extraction of flies) and evolved replicate samples and their respective generation 0 expansion sample (‘Gen. 0->n’). Grey points indicate individual replicate samples, and black diamonds the averaged value across replicates. F_ST_ differentiation from generation 0 samples increased monotonically as function of collection time (F = 172.2; p-value < 0.01). (B-C) MDS of F_ST_ values computed pairwise across all expansion samples, colored according to (B) replicate cage or (C) collection generation. (D) Average Pearson Correlation across replicates (+/- standard deviation) between sample expansion generation and distance along a one-dimensional axis constructed using F_ST_ MDS coordinates for early expansion (generation 0-4) samples (Fig. 2C). Purple points and error bars correspond to observed values (mean +/- standard deviation across cages), while grey points and error bars correspond to values derived from N = 100 permutations. (E) Trajectories of rising alleles at SNPs identified via GLM (FDR < 0.01 and effect size > 2%) across all nine generations of expansion. (F) Genomic distribution of SNPs depicted in (E). The coordinates of the major cosmopolitan inversions segregating in our founding reference panel at greater than 4% frequency in our founding strains are depicted above the x-axis. (G) Leave-one-out cross-validation to infer replicate-specific parallelism of adaptation to sustained population expansion. Portrayed is the median shift of the rising allele for sets of target SNPs (points) identified via GLM (FDR < 0.01 and effect size > 2%) in 3 of 4 cages, relative to medians derived from a matched control SNP set (X’s), in the left-out replicate. Target SNP medians are colored red if the distribution of phased allele frequency shifts was significantly greater than that of matched control sites (two-tailed t-test, FDR < 0.05).

To more rigorously quantify the systematic movement of samples through time observed in the two-dimensional MDS space, we translated coordinates for each cage such that the centroid of all generation 0 samples was centered at the origin. We then used these translated points to fit a simple linear regression model to samples from expansion generations 0-4 (Methods; Fig. S5). The resulting axis represents the primary axis of variation in the 2D plane during early expansion for each replicate. We hypothesized that if sustained, directional selection imposed by population expansion was a primary driver of patterns of genomic variation, samples from the remaining expansion generations (generations 6-9) would continue to proceed along the established axis of variation in the same direction. Indeed, projection of late expansion samples onto this axis of variation indicated a significant correlation of sample collection generation and distance along the axis (Fig 2D.; significance of correlation determined via permutations). When segregating this analysis by chromosomal arm, however, this parallelism was only evident on chromosomal arms 2L, 3L, and 3R ; while correlations derived from SNPs on 2R and X were indistinguishable from permuted values (Fig. 2D). We re-sampled SNPs across the genome iteratively to match the number present on each chromosomal arm to confirm that the variation in parallelism was not a technical artifact. In each case, our sub- sampling yielded correlations that matched the genome-wide trend, indicating that differences observed among chromosomal arms (Fig. 2D) reflect systematic differences in the behavior of loci across the genome (Fig. S6, Table S2).

To determine the distribution of SNPs (N = 1.7 M SNPs) contributing to the parallel patterns of divergence across cages, a generalized linear model was fit to allele frequencies from all expansion samples (generation 0-9). This model assessed the significance of the linear relationship between allele frequency and generation of sampling across all cages, using a quasibinomial error model to reduce false positive associations (15,16,34). While an association between allele frequency and sampling timepoint within a single cage may represent either drift or selection, a significant parallel association across all four replicate cages indicates allele frequency trajectories that are both predictable over time and parallel across populations. In this case, selection (or linked selection) is the more parsimonious explanation. A total of 389,588 SNPs (22.9 % of all sites), across all chromosomal arms, showed significant parallelism after multiple testing correction (Benjamini-Hochberg false discovery rate <0.01 and allele frequency change > 2%). These SNPs showed systematic, directional selection across replicates throughout expansion (Fig. 2E; Fig. S7). Furthermore, the genomic distribution of these SNPs spanned all five chromosomal arms and were located both within and outside the breakpoints of five major cosmopolitan inversions that segregated at appreciable frequency (>4%) in our inbred reference panel (Fig. 2F).

We next used our GLM signal to infer the minim number non-overlapping, putatively unlinked loci, underpinning patterns of adaptation throughout expansion (ref; Methods). This resulted in the identification of 250 total loci, ranging in size from 30 kb to 305 kb and located across all chromosomal arms (Supplementary File X). Allele frequency trajectories of the most significant SNPs within each clusters indicate strongly directional selection throughout expansion, with median selection coefficients of 7% per generation (Fig S8). The number of unlinked loci and the inferred strength of selection is on par with similar estimates obtained from outdoor mesocosom and wild population sampling (14–16) While determining the underlying causal regions (which could be either SNPs or larger structural variants) within these unlinked loci requires further investigation, this analysis at least suggests adaptation in response to increasing population densities to be underpinned by a fairly large number of independent variants (i.e. on the order of hundreds), each containing alleles exhibiting relatively large selection coefficients. We further note that while long-range patterns of linkage could theoretically induce non-independent patterns of allele frequency change among loci on the same chromosomal arm, we have previously shown that how this method, which identifies independent loci based on patterns of physical linkage, produces sets of loci with independent allele frequency movement throughout evolution (14).

Finally, we validated the parallelism inferred by GLM by implementing a leave-one-out cross validation. Specifically, we iteratively identified sets of parallel SNPs across expansion samples using a GLM fit to allele frequency data from three of the four replicate cages. We then quantified the frequency shifts of the rising allele at significant SNPs (FDR < 0.01, allele frequency shift > 2 %) in the left-out cage. In each iteration of this analysis, the left-out cage exhibited a magnitude of allele frequency change that exceeded the background allele frequency change (quantified using matched control SNPs; see Methods), and in a direction of change parallel to that observed in the other three cages (Fig 2G). The genome-wide median shifts of target SNPs across replicates ranged between 5 and 7.5%, indicating that these parallel patterns of adaptation were underpinned by strong selection, on the order of 10-20% per generation. While the per-chromosome Fst and MDS analyses described above, which focus on signal averaged across all segregating sites, suggest that non-parallel idiosyncratic movement may be the dominant force on certain chromosomes, our GLM analysis parallel SNPs can still be discovered on every chromosomal arm.

### Genomic evidence of trade-offs induced by fluctuating selection across expansion and truncation

The parallel frequency shifts observed throughout expansion may be the product of three, not mutually exclusive, evolutionary dynamics: (1) directional selection in response to sustained fecundity selection, (2) adaptation to the lab environment, and/or (3) the purging of recessive deleterious alleles (i.e., negative selection) in outbred population. To disentangle these various dynamics and, in turn, identify the presence of fitness-tradeoffs at putatively selected alleles, we leveraged our samples collected throughout truncation selection. Alleles identified during expansion that were a product of consistent lab selection and/or negative selection against unconditionally deleterious recessives should continue to show systematic, directional change throughout truncation (as their effect is not conditional on the specific treatment). However, alleles with treatment-specific behavior during expansion and truncation (i.e., moving in the opposite direction) represent those that likely underpin trade-offs between fecundity and stress tolerance selection.

We tested for evidence of context-specific behavior and trade-offs of genome-wide SNPs by re-conducting our MDS analysis of pairwise divergence values, this time including all samples collected throughout expansion and truncation. If the dominant direction of allele frequency change was sustained across both expansion and truncation (indicating sustained lab selection or purging of deleterious mutations), truncation samples would be ordered from early to late in a parallel manner as the expansion samples (Fig 2B). Instead, we observed that samples taken during truncation (initiated at expansion generation 5) shifted back towards earlier expansion samples, potentially suggesting a genome-wide reversion of allele frequencies (Fig. 3A; Fig. S9). We quantified these trends as above, translating the MDS coordinates of each truncation sample such that the centroid of all hour 0 samples was centered at the origin and then projecting them onto a the single axis linear regression model derived from early expansion points (Fig. 2D; Fig. S5). As hypothesized, correlations between collection time and distance along this axis were, genome-wide, significantly negative (Fig. 3B; Table S1). Segregating this analysis by chromosomal arm yielded more nuanced dynamics whereby 3L, 3R, and X yielded evidence of anti-parallel/reversions in allele frequencies, while 2L and 2R exhibited correlations that were insignificant relative to our permuted distributions (Fig. 3B). Again, as above, we validated that these differences among chromosomal arms were not simply a reflection of variation in number of SNPs available for measurement and reflected systematic differences in allele frequency dynamics across the genome (Fig. S10).

**Figure 3.**
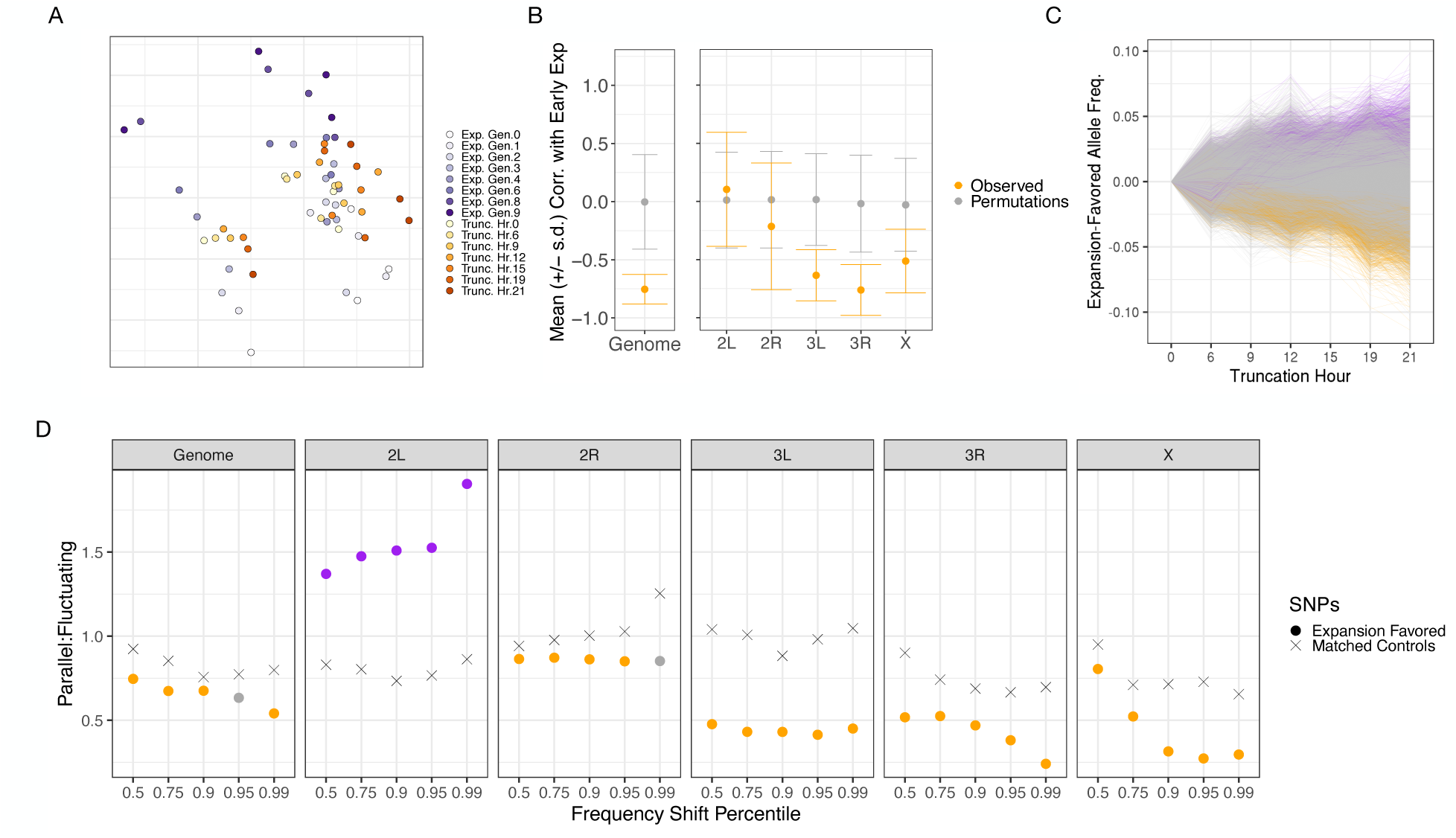
Genome-wide signal of fitness trade-offs between population expansion and truncation. (A) MDS of pairwise F_ST_ values across all samples collected throughout expansion (purple-hue points) and truncation (orange-hue points), shaded according to collection time point (darker hues indicate later expansion or truncation sampling generation/hour. (B) Average Pearson correlation across replicates (+/- standard deviation) between sample expansion generation and distance along a one-dimensional axis constructed using F_ST_ MDS coordinates for early expansion (generation 0-4) samples (Fig. 3A). Orange points and error bars correspond to observed values (mean +/- standard deviation across cages), while grey points and error bars correspond to values derived from N = 100 permutations. (C) Trajectories of SNPs identified via GLM regression across replicates throughout expansion, measured during truncation. Trajectories are colored for SNPs that exhibited evidence of systematic allele frequency changes across replicates during truncation (GLM FDR < 0.05 and allele frequency change > 1%) and displayed sustained directional (purple) or fluctuating selection (orange). (D) Ratio of parallel to fluctuating dynamics for expansion-favored alleles during truncation, segregated by chromosomal arm and percentile of truncation allele frequency shift. Points are further colored if the ratio of parallel to fluctuating behavior differed significantly from that expected based on a matched control set (χ^2^ test P-value < 0.05).

We garnered further evidence of trade-offs across fecundity and desiccation selection using allele frequency trajectories. Specifically, we selected those SNPs with evidence of systematic movement across replicates throughout expansion (GLM FDR < 0.01 and allele frequency shift > 2%) and measured the average frequency shifts of the rising allele across cages throughout truncation. If patterns of fluctuating selection and antagonistic pleiotropy dominated the data, we expected trajectories to be enriched for SNPs moving in the opposite (negative) direction during truncation. Indeed, across all expansion-identified SNPs, the ratio was significantly skewed towards those with exhibiting fluctuating selection, as opposed to sustained directional selection, across expansion and truncation selection (χ^2^ = 1123.6; P-value < 0.001). These trajectories are depicted in Figure 3C, in which colored trajectories indicate those expansion-identified SNPs with additional, independent evidence of systematic movement across cages during truncation (GLM FDR < 0.05 and effect size > 1%): purple for those SNPs with consistent directional selection across selection regimes, and orange for those with anti- parallel movement across regimes. As expected, this subset of SNPs (*N =* 24,814) exhibited even greater evidence of fluctuating selection and were 3.5 times more likely to fluctuate directions across treatments than exhibit sustained directional selection (19,158 relative to 5,656 SNPs, respectively). We assessed how consistent this dynamic was across the genome by quantifying the ratio of parallel vs. fluctuating selection genome-wide, and separately for each chromosomal arm. As expected based on the F_ST_ MDS analysis described above, there was strong evidence of enrichment for fluctuating selection on chromosomal arms 3L, 3R, and X. However, this analysis revealed an enrichment of 2L SNPs with sustained parallel selection across selection regimes, which was an undetected in our F_ST_ MDS analysis. We explored this dynamic further, using the underlying allele frequency shift distributions to compute empirical cumulative distribution functions, separately for alleles with sustained directional selection and for those with evidence of fluctuating selection (See Supp. Mat.). This verified a genome-wide enrichment of alleles exhibiting fluctuating selection (driven by chromosomal arms 3L, 3R, and X), and evidence of sustained directional selection on 2L (Fig S11). This signal of fluctuating selection was further corroborated by an assessment of the behavior of unlinked loci identified during reproduction selection throughout truncation (Supplementary Data File 2, Figure S12). In sum, we find overwhelming evidence for the existence of pervasive, genome-wide trade-offs between fecundity and desiccation selection, as well as some evidence of sustained directional selection due to either adaptation to the lab environment or purging of the unconditionally deleterious alleles. It is important to note the patterns of fluctuating selection quantified here cannot be a spurious artifact driven by regression to the mean, as the allele frequency data from which the trajectories were computed are entirely independent across expansion and truncation (i.e., have no overlapping time points).

### One and two locus simulations of linkage, selection, and antagonistic pleiotropy

We explored whether patterns of linkage between loci, antagonistic pleiotropy at a single SNP, and/or haplotype-level antagonistic pleiotropy could generate the patterns of allele frequency change observed in our data using Wrigh-Fisher simulations in SLiM (35). Briefly, we first simulated a scenario in which a single SNP had an advantageous effect on fitness during expansion, which was countered by a disadvantageous effect during truncation. As expected, this produced frequency trajectories across reproduction and stress tolerance selection that recapitulated the patterns observed in our empirical data (Fig S13).

Next, we focused on a series of two-locus scenarios in which SNP ‘A’ was directionally favored during expansion with neutral fitness effects during truncation, while SNP ‘B’ had neutral fitness effects during expansion and then came under directional selection during truncation. We then computed fitness values (empirical selection coefficients) during the first generation of truncation (which represents the single generation (no-recombination) of stress tolerance selection in in our experiment) to infer how the relationship between the two SNPs may influence their behavior across phases. Under scenarios in which the two loci were unlinked, their frequency trajectories were largely independent across phases: SNP ‘A’ increased systematically during expansion with neutral dynamics during truncation, while SNP ‘B’ exhibited neutral dynamics during expansion and then systematic increases during truncation (Fig S14-S16). Under a scenario of high linkage and a net ‘attraction’ between the favored allele at each SNP (i.e., the favored allele at SNP ‘A’ was non-independently assorted with the favored allele at SNP ‘B’), trajectories were largely correlated. For example, SNP ‘A’ increased in frequency systematically across both expansion and truncation, causing empirical selection coefficients to indicate a selective advantage during truncation even though the locus was in fact neutral during this phase (Fig S14; Fig S17-18). Alternatively, under a scenario of high linkage and net ‘repulsion’ between the favored alleles at each SNP (i.e., the favored allele at SNP ‘A’ was non-independently assorted with the disadvantageous allele at SNP ‘B’), SNP ‘A’ exhibited a trajectory of fluctuating selection whereby it systematically increased in frequency during expansion and then declined during truncation (Fig S14; Fig S19-20). This scenario thus represents a form of ‘haplotype-level’ antagonistic pleiotropy driven by the orientation of causal alleles and the underlying haplotype structure in the population. Finally, under a scenario in which the attracted and repulsed states of the selected alleles occur at equal probability (i.e., random assortment of alleles between loci), the net-effect of possible linkage patterns produces allele frequency trajectories and empirical fitness values that recapitulate the dynamics observed in the ‘unlinked’ scenario (Fig S14; Fig S21-22).

As we have no prior knowledge of a non-independent assortment of selected alleles in our focal population, the final simulated scenario (independent assortment of advantageous alleles between locus ‘A’ and ‘B’) likely represents the most plausible dynamic occurring in our experiment, should two linked SNPs be under differential selection across selection regimes.

Therefore, within the confines of the current data (i.e., not yet knowing the underlying causal loci), we conservatively interpret our observations of reversions in allele frequency trajectories as antagonistic pleiotropy at a single SNP, though recognize the possibility that it could in fact represent antagonistic pleiotropy at the haplotype-level. Understanding how such haplotype structure could be maintained over long time periods, via either strong selection for the ‘repulsed’ linkage between selected alleles or some form of recombination suppression (e.g., an inversion), will ultimately hinge upon resolving the causal loci in this system.

### The role of inversions in reproduction/stress-tolerance trade-offs

Theoretical, and emerging empirical, research suggests a dominant role of chromosomal inversions in adaptation in natural populations (36). Accordingly, we quantified whether five major cosmopolitan inversions in *D. melanogaster* (*In(2L)t*, *In(2R)NS*, *In(3R)K*, *In(3R)P*, *In(3R)Mo*), which all occurred at starting frequency > 4% in our inbred reference panel, exhibited dynamics consistent with adaptation and, if so, trade-offs between population expansion and truncation (Supplementary Data File 3). Generalized linear regression of inversion frequencies yielded evidence that all inversions shifted systematically across replicates during expansion, with magnitude of frequency shifts ranging between 1 and 7*%* (Fig. 4A-E; Table S2). *In(3R)K* and *In(3R)P* were the only two inversions that exhibited evidence of trade-offs across selection regimes, whereby each inversion was systematically favored during expansion and then became selected against during truncation (Fig. 4A-B).

**Figure 4.**
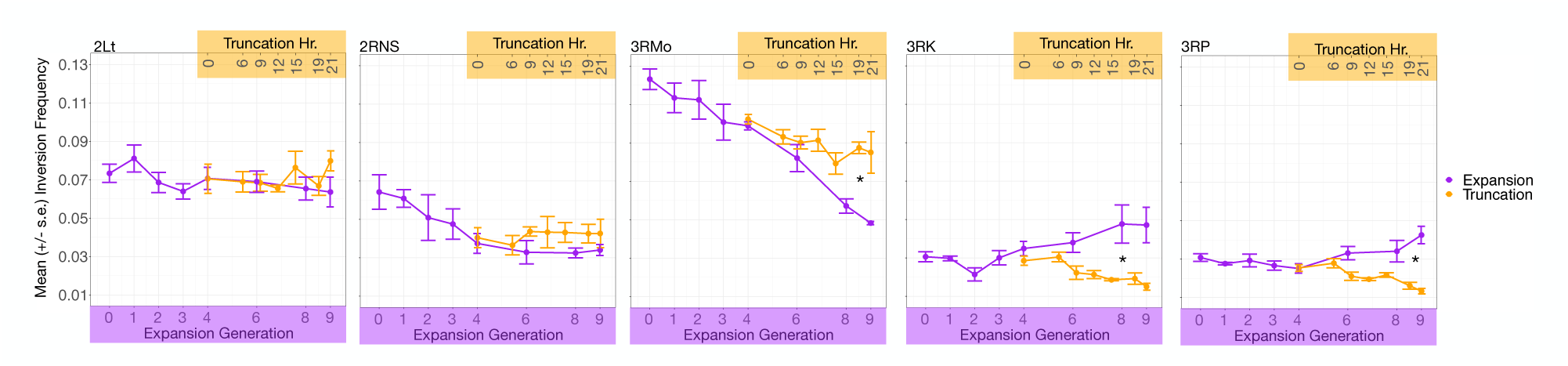
Role of inversions in reproduction and stress-tolerance trade-offs. Mean inversion frequency changes (+/- s.e.) across expansion generations (purple trajectories) and truncation hours (orange trajectories). Asterisks indicate inversions with evidence of fluctuating selection across expansion and truncation.

Given the appreciable effect of inversions on patterns of adaptation throughout the experiment, we next explored the extent to which our inference of trade-offs was solely a function of variation within, or in tight linkage between, these structural variants. Specifically, we reconducted F_ST_-based MDS analysis on a subset of SNPs, excluding all those within, or 100 kb away from, inversion breakpoints. Through this, we observed that the signature of trade-offs was maintained on 3L and X, but eliminated on 3R. Still, the genome-wide signal of this analysis continued to indicate parallel movement across early and late-expansion samples, and that a signal of fluctuating selection dominates patterns of genomic variation between expansion and truncation (Fig. S23). We validated that this trend was not simply a function of the reduced set of SNPs available for analysis, relative to the genome-wide panel (Fig. S24).

### The emergence of tradeoffs in response to natural environmental fluctuations

We quantified whether alleles identified via selection under sustained population expansion in a laboratory setting can predict patterns of adaption and trade-offs in an outdoor environment, where populations adapt both to changes in population density as well as a suite of additional abiotic variables. Specifically, we leveraged data from an independent study year when we monitored patterns of genomic variation in a genetically diverse population that was split into a series of large, replicate cages maintained in both a controlled, indoor laboratory (N = 10 replicate populations), as well as outdoor mesocosms exposed to natural environmental fluctuations (N = 12 replicate populations; data previously reported in Bitter et al. 2024) (Fig. S25). As with our indoor expansion/truncation experiment described above, the population used in this experiment was derived via outbreeding an inbred reference panel, which in this case was originally collected from Linvilla Orchards, Media, PA (Supplementary Data File 4). The replicates in both environments evolved with overlapping generations, under constant food conditions, for a period of four months (see Methods). This induced rapid population expansion in each environment until a peak/stabilization of density was observed, suggesting the initiation of density control (Fig. S25; Table S7).

We first quantified the extent to which evolution in the outdoor environment is driven by the selective pressures solely associated with increasing population density (e.g., selection for increased fecundity, faster developmental rate). Specifically, we identified alleles systematically favored throughout concurrent sampling of the indoor and outdoor cages using a GLM (see Methods), and found far greater overlap than expected based on matched control SNPs (Fig. 5A). Interestingly, we failed to detect such significant statistical overlap using the set of loci identified during the reproduction selection experiment described above, which used a different inbred reference panel (see Methods). Next, we asked whether the dominant direction of selection was parallel across the indoor and outdoor mesocosm by quantifying frequency shifts in the outdoor replicates at the rising allele for each SNP identified within the indoor environment. We observed that the magnitude of allele frequency shifts was significantly greater than background allele frequency movement, and that the dominant direction of selection was conserved between environments (indicated via positive median shifts depicted in Fig. 5B; Table S6).

**Figure 5.**
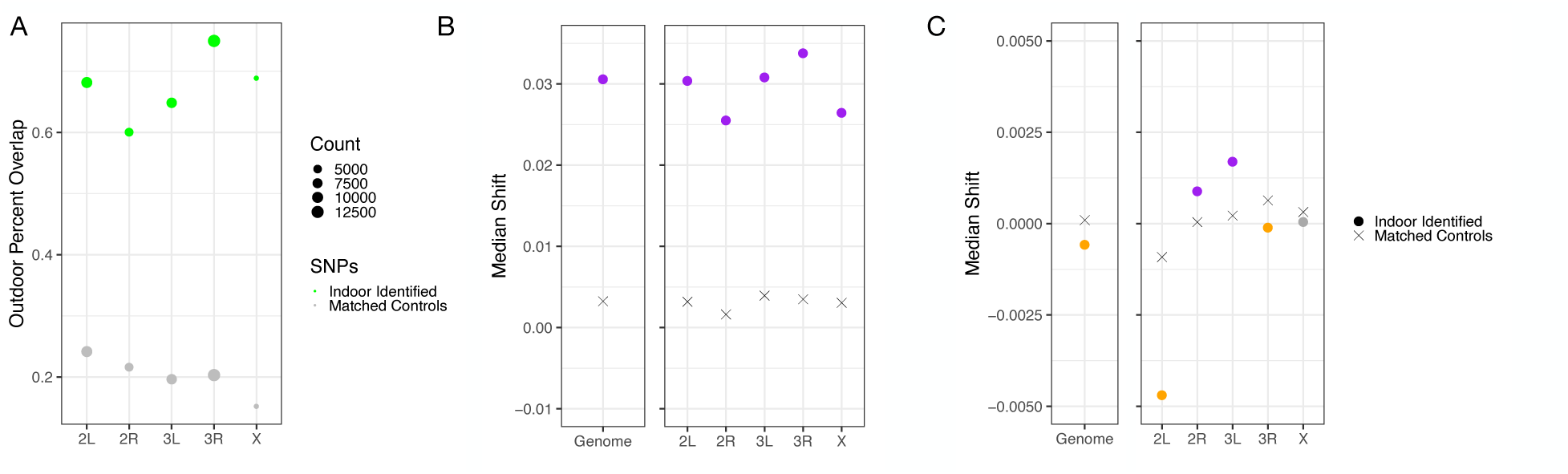
Selection on reproduction under controlled, lab-based conditions predicts patterns of adaptation and trade-offs in outdoor mesocosms. (A) Observed vs. expected overlap in SNPs with systematic allele frequency movement (GLM FDR < 0.01; allele frequency change > 2%) in a paired indoor and outdoor experimental evolution study. (B-C) Median shift of alleles identified during expansion in an indoor environment, quantified during population expansion (B) and collapse (C) in an outdoor mesocosm. Colored circles correspond to indoor-identified SNPs with allele frequency shift distributions that were either significantly greater (orange) or less (purple) than that observed for matched control SNPs (X’s) (two-tailed t-test FDR < 0.05).

Finally, we tested whether the pervasive trade-offs we inferred from the previous experiment and results described above (Fig. 1-4) manifested in the outdoor mesocosms. Specifically, we leveraged an additional month of sampling in the outdoor mesocosms, during which time a population decline and ultimate collapse was observed as winter and the deterioration of abiotic conditions progressed (14). We hypothesized that, should those alleles identified in the indoor environment underpin antagonistic pleiotropy and trade-offs for fitness-relevant variation, allele frequencies should reverse in direction as the outdoor mesocosms collapsed. Indeed, we found a significant genome-wide reversion in allele frequencies in the outdoor cages throughout this period, a dynamic most primarily driven by patterns of variation on 2L and 3R (Fig. 5C; Table S6). This shift in the dominant direction of indoor-identified SNPs indicates that their signatures of selection are not simply driven by the purging of deleterious recessives in our reference panel, but rather exhibit context-dependent fitness effects across population expansion and truncation.

## Discussion

Here, we used experimental evolution in large, genetically diverse populations of *D. melanagoster* to quantify genome-wide evidence of fitness trade-offs in response to selection elicited in the context of ecologically realistic population boom and bust dynamics. We first manipulated reproduction selection via nine generations of sustained population expansion in the absence of density regulation mechanisms. This process induced parallel, genome-wide shifts in allele frequencies across four, independent replicate populations. The strong selection coefficients quantified throughout this period (∼7% per generation) are comparable to those quantified during sampling between summer and fall in wild populations and outdoor mesocosms (13–16). Thus, while it has been previously speculated that such patterns of evolution of *D. melanogaster* across seasons may be dominated by adaptation to shifting temperatures (e.g., Boulétreau-Merle, Fouillet, and Terrier 1987), our lab-based manipulation here provides direct evidence that the sole impact of shifting population densities can drive rapid adaptation over the course of several generations.

By imposing a bout of truncation selection midway through population expansion, during which time individual survival and associated patterns of allele frequency change revealed differences in relative stress tolerance (e.g., desiccation resistance) among genotypes, we aimed to explore evidence of fitness-tradeoffs in the system. We quantified a pervasive, genome-wide signal of fluctuating selection, whereby patterns of allele frequency movement reversed direction across selection regimes. These patterns are indicative of antagonistic pleiotropy, whereby selected alleles conveying advantageous trait values for the phenotypes favored during reproduction selection convey disadvantageous trait values for the suite of phenotypes favored during truncation (7,37). While this dynamic was *a priori* expected based on the negative genetic correlations between traits likely under selection in our experiment (10,29–31), few data exist directly observing the manifestation of such trade-offs, particularly over the ecologically-relevant timescales assayed in our experiment. The key implication of this antagonistically pleiotropic behavior is that this process may, in effect, maintain variation in the population (7,8). Specifically, mutations with unconditionally advantageous effects will ultimately be driven to fixation, while those with context-specific behavior can in principle be maintained for longer periods of time in the presence of fluctuating selective pressures (38,39). The key challenge for future work will be to discern whether these dynamics indeed balance alleles over long time-periods, which will ultimately be aided by refining the causal loci driving the patterns observed here and observation across multiple bouts of fluctuating selection.

The genomic architecture of trade-offs was not dominated by a single, large-effect locus, nor were antagonistically pleiotropic alleles distributed evenly across chromosomal arms. Rather, we quantified a dominant signal of fluctuating selection across reproduction and stress-tolerance selection on chromosomal arms 3L, 3R, and X, and extensively showed that this signal was not spurious due to variation in the number of SNPs available for analysis across arms. We further showed that the trade-offs quantified on 3R were likely underpinned by inversions segregating in our experimental population. Inversions are a common form of structural variation both across species and within populations and have long been recognized as important for adaptive evolution and, more recently, in trade-offs (21). Mechanistically, suppressed recombination within the breakpoints of relatively young inversions can give rise to the accumulation of putatively adaptive alleles (40–43), and over long periods maintain allelic structure within populations that generate haplotype-level, antagonistically pleiotropic fitness effects (see *One and two locus simulations of linkage, selection, and antagonistic pleiotropy,* and Fig. S14). In natural populations of *D. melanogaster*, inversions have indeed been implicated in adaptation to spatially and temporally varying selection pressures, which may, in part, be underpinned by the selective pressures manipulated in our lab-based study (15,44–46). We add to this growing body of research, here reporting evidence for a role of each of five major chromosomal inversions during reproduction selection, with two of these inversions, *In(3R)P*, and *In(3R)K*, showing a behavior consistent with trade-offs between reproduction and stress-tolerance selection. In congruence with this result, previous experimental work has demonstrated that inverted karyotypes of *In(3R)P* are shorter-lived and less stress resistant than non-inverted karyotypes, as would be predicted based on the frequency trajectories of this inversion in our experiment (47). Still, resolving the functional relevance of each of these major inversions is nascent and provides a fruitful avenue of future research.

It is noteworthy that the trade-offs quantified here manifested during a single bout of truncation selection, which proceeded over the course of just 24 hours. This illustrates how finely adaptation via standing variation can track shifting environmental conditions, and that selection can be detected on temporal scales shorter than a single generation. Indeed, mounting research has shown how both anomalous (e.g., heat wave, hurricane, or poaching) and non-anomalous (e.g., weekly to seasonal shifts in abiotic and biotic conditions) environmental perturbations can elicit patterns of adaptation on similar timescales in natural populations (13–16,46,48–51). This growing body of research increasingly suggests that the selection coefficients underpinning the standing, functional variation in natural populations may be substantially larger than previously recognized, and ultimately augment adaptation to the accelerated shifts in mean environments associated with global climate change.

Our paired indoor cage and outdoor mesocosm experiment demonstrated how selection induced by population expansion under controlled, indoor conditions can identify alleles that are relevant to adaptation and trade-offs in response to natural environmental fluctuations. The conditional benefit of the indoor-identified loci in the outdoor mesocosm (i.e., their genome-wide advantageousness during population expansion and genome-wide disadvantageousness during truncation; Fig. 5) indicates that these the shared signatures of selection across environments do not simply represent a set of loci under negative selection (e.g., purging of deleterious recessives), but rather subject to some form of balancing selection imposed by the temporally fluctuating environment. In turn, this finding indicates that the repeated signals of fluctuating selection quantified in *D. melanogaster*, both across populations and through time (13–16), may be underpinned by fundamental life-history trade-offs that emerge as a result of shifts in the selective environment induced by population boom-bust demographic dynamics. These boom-bust demographic dynamics are a generic feature of populations across taxa, and this phenomenon may thus be more widespread than previously recognized and provide a key instance of the interplay of ecological and evolutionary forces in natural populations (52–55).

Furthermore, given the dramatic differences in indoor and outdoor environments, the shared allele frequency patterns observed here suggest that core alleles may underpin adaptation during generic boom and bust cycles, regardless of specific abiotic condition. This bolsters the hypothesis that the predictability of adaptation in this system is augmented by the nature of the underlying genetic variation: loci responding to rapidly fluctuating selection are not specific in their response to particular abiotic parameters, but rather respond more coarsely to generic and repeatable features to the selective environments, in this case the population density fluctuations that occur yearly in the system. Such a ‘coarse-graining’ of the architecture of the adaptive response would position life-history trade-offs and fluctuating selection as a key force in maintaining variation in natural populations. Ultimately, fine-mapping the specific causal loci underpinning these trade-offs and signatures of fluctuating selection is a key area of future research that, as the data presented here suggest, may even be feasible within the context of highly replicated indoor experiments. Finally, we note the possibility that a unique set of loci become fitness-relevant under specific abiotic contexts, our goal here was to simply query whether a common set of loci underpinned trade-offs and responded generically to fluctuating selection across abiotic contexts, for which we found substantial evidence.

It is notable that we observed a putative difference in the architecture of trade-offs between the indoor population expansion/truncation selection experiment, and the paired indoor-outdoor mesocosm study. For example, chromosomal arm 2L displayed evidence of sustained selection across reproduction and stress tolerance selection in the first experiment, but exhibited strong evidence of antagonistic pleiotropy between population expansion and collapse during our outdoor mesocosm experiment. Furthermore, we did not detect any shared enrichment of SNPs with evidence of linked selection across these experiments. While there were distinct methodological and environmental differences that may have played a role in this discordance, a more salient possibility is the use of a different set of inbred lines to generate the outbred mapping population for each experiment. As our analyses ultimately identify sets of SNPs in tight linkage to an underlying causal locus, differences in patterns of linkage across mapping populations could, in effect, lead to dramatic differences in the relative effect size of marker alleles across studies. This process is analogous to the oftentimes poor portability of association studies across human populations (56,57), and is an important consideration for future research using experimental evolution approaches to characterize the architecture of complex trait adaptation.

In conclusion, the set of experiments conducted here demonstrate how adaptation associated with population boom-bust ecology impose selection on a core set of loci, eliciting trade-offs and driving predictable evolutionary dynamics in populations of *D. melanogaster*. This dynamic, in turn, likely acts as a key force maintaining variation in the species, and potentially a broader range of taxa experiencing similar boom-bust demographic dynamics. Finally, this study demonstrates how well-resolved time-series genomic data can reveal the presence of, and genomic architecture underlying, fitness trade-offs.

## Methods

### Population construction and replicate cage seeding for expansion/truncation selection experiment

We constructed a genetically diverse founder population via outbreeding 145 lines of the Drosophila melanogaster Genetic Reference Panel (DGRP; Mackay et al. 2012). Ten mated females per DGRP inbred line, each of the same age cohort and from density-controlled line cultures, were pooled into a single 0.3m x 0.6m x 0.3m cage (P0 generation). Over the course of 4d, eggs were collected in 64 culture bottles (P1 generation) using cornmeal molasses medium (16 bottles added per day, then capped, removed, and 16 new bottles added); the 64 bottles were then randomly assigned to 1 of the 4 replicate cages; from this point forward, all replicates were cultured independently. Once flies eclosed, they were again released into 4 replicate 0.3m x 0.7m x 0.3m cages, and eggs were collected over 24h on two culture trays (0.5m x 0.3m x 0.1m; 1L of Drosophila cornmeal molasses medium per tray) per replicate. Once these flies eclosed (the first true F1 generation), the flies were released into 4 replicate, medium sized cages (0.6m x 0.6m x 1.2m) and allowed to oviposit on 4 trays of culture medium over 24h (embryos were the F2 generation). The F2 embryos from these trays were sealed and collected, and the adults discarded. Once the F2 flies eclosed, they were then released into large, experimental cages (3 m^3^) and given 8 trays of media for oviposition over 24h.

### Population expansion/truncation selection in large, indoor cages

Culturing within the large, experimental cages proceeded via discrete generations, throughout which the amount of food was doubled every generation (starting at 8 trays for the F2). By doubling food every generation we generated sustained population expansion, thereby continually selecting for the fecundity-associated traits that are advantageous in wild populations when resources are abundant, most likely reproductive output and increased developmental rate. Hereafter, we refer to collection timepoints in accordance with the number of generations since release into large cages (the first 8 tray stage), which was two generations from the actual founding of the experimental populations (i.e., the samples labeled “generation 0” are F2’s, “generation 1” are F3’s, and so forth). Once flies had laid eggs for a period of 24h, the adults were removed from each cage and their volume measured to estimate census size.

The remaining embryos (i.e., subsequent generation of flies) were left to develop and eclose within the same replicate cage. Across replicates, the population sizes rapidly expanded across generations, a dynamic selecting for individuals exhibiting increased reproduction (e.g., fecundity and developmental rate) (Fig. 1).

We maintained food doubling until generation 4 at which point, due to logistical constraints, it was not possible to continue the doubling of population size per cage beyond the 64 tray per cage generation. In order to continue the selection regime of population expansion (and selection for early fecundity, fast developmental rate), we instituted a random dilution followed by re-expansion. Specifically, the fifth generation was founded with embryos collected over 24h on two food trays (containing eggs of approximately 20,000 flies), from the from the fourth generation. These generation 5 flies were allowed to develop and eclose in the large indoor cages, at which point 8 trays of medium were added to each replicate cage, and oviposition was carried out for 24h and after which adults were discarded. Once the sixth generation of flies eclosed, they were allowed to oviposit for 24h on 2 food trays. These generation 7 embryos were then allowed to develop, eclose, and expanded out to 8 trays for the eighth generation. The generation 8 flies were allowed to oviposit over 24h on 8 trays, consistent with the previous generation (thus each generation of flies resulted from egg laying over a period of 24h from the preceding generation). We collected a random sample of 100 male and 100 female flies in generations 0, 1, 2, 3, 4, 6, 8, and 9, and pooled sexes separately for later whole genome shotgun sequencing (see below). We took additional samples of each replicate at generations 6 and 8 to use as biological replicates (i.e., same replicate/generation, different set of 100 flies) to quantify noise in our allele frequency estimates (see below). Census estimates (based on the total volume of dead flies) of adult flies within each cage were conducted on generations 1-4. Calibration of census estimates was conducted by counting the number of flies (desiccated and dried) in 1cm^3^ and then measuring the volume in a given sample.

Concurrent to the generation 4 dilution and re-expansion described above, we induced a bout of truncation selection that segregated flies based on stress-tolerance, most likely desiccation resistance as water availability is expected to drive mortality at a much faster rate than starvation (33). Specifically, after collecting generation 5 embryos for continued expansion, we retained the adult flies within their respective cages, removed all food and water, and sampled the surviving adults at various timepoints by direct aspiration using vacuums until all flies had died. The live samples were immediately sorted by sex and timepoint and preserved in ethanol (as with expansion samples, 100 male and 100 female flies were isolated at each collection time point for whole genome shotgun sequencing). We continued this process until there were an insufficient number of flies (i.e., < 100 individuals) for sampling, resulting in samples for DNA analysis collected at hours 0, 6, 9, 12, 15, 19, and 21. Samples collected throughout this process were expected to reveal changes in the relative frequency of genotypes of differing degrees of stress-tolerance, traits expected to trade-off with the reproductive traits favored during the nine generations of expansion selection (10,29).

### Pooled genomic sequencing and allele frequency estimation of expansion/truncation selection experiment

Genomic DNA from pools of 100 male and 100 female flies were extracted and sequenced in two rounds. First, multiplexed libraries were created for 56 expansion samples using Illumina Nextera DNA Prep with e-gel size selection and i7 indexing. Barcoded fragments were mixed and loaded evenly into 4 lanes, then sequenced with 100 bp paired-end reads on an Illumina HiSeq2000 sequencer, with target coverage of 10x per sample. After QC, resulting per/sample coverage was quite variable, and 26 samples from this round with coverage <2x were later re-sequenced with 150-bp dual-indexed reads on an Illumina HiSeq4000 sequencer.

Reads from the same sample were merged after all QC and mapping steps. A separate round of sequencing was conducted for the remaining 96 samples, including remaining expansion and truncation samples. These samples were sequenced with 150-bp paired-end dual-index reads on a NextSeq550 high output machine, with a target coverage 7x per sample. All samples were demultiplexed, adapter sequences were trimmed, and reads with any 3’ bases with quality score < 20 or trim length <18 were discarded. Overlapping forward and reverse reads were subsequently assembled and reads were mapped separately to the *D.mel* v5.39 reference genome using bwa and default parameters (58). Aligned reads were deduplicated using Picard tools (http://broadinstitute.github.io/picard/), and all reads were re-aligned around indels using GATK v4 IndelRealigner (https://gatk.broadinstitute.org/).

We used the founder line genome sequencing data to compute haplotype-informed allele frequency estimates at 2.7 M, previously identified, segregating sites using a local inference method and pipeline developed and described by (59,60). We conducted haplotype inference in window sizes that varied proportionally to the length of un-recombined haplotype blocks expected as a function of the estimated number of generations since the construction of the outbred population. We have previously provided extensive validation that our haplotype-informed allele frequencies are replicable across different sets of subsampled reads from the same sample, and produce an accuracy of allele frequency estimates that is comparable to deep sequencing and standard methods for pooled, allele frequency estimation (14,16,60). We filtered our allele frequencies to only include those sites with an average minor allele > 0.02 across all samples, resulting in 1.7M SNPs for analysis. Finally, we averaged technical replicate allele frequencies (i.e. the male and female pool) for each replicate and collection time point, resulting in 4 cages per time point, across 8 expansion generations and 7 truncation time points (60 total samples).

### Identifying parallel patterns of allele frequency change and genome-wide evidence of trade-offs

Statistical analysis of allele frequency data was conducted using R v. 3.5.6. We first explored patterns of genomic divergence between samples as average F_ST_ across all segregating sites, which provides a metric of allele frequency divergence between samples that is normalized by the starting frequency of the allele. We compared F_ST_ between biological replicates (same replicate cage/collection time point, different pool/extraction) to those values obtained for each replicate cage between its generation 0 sample and all subsequent generation expansion samples. We quantified whether evolutionary divergence increased throughout the course of expansion, and at what point it exceeded the sources of biological noise impacting our allele frequency estimates, using a linear regression.

We next explored whether increasing differentiation through time was underpinned by parallel allele frequency shifts across replicates. Specifically, we used our pairwise divergence values (i.e., F_ST_ between samples) to create multi-dimensional scaling (MDS) plots, in which divergence between samples is represented as distance between points in a 2-D plane (*cmdscale* function in *stats* package R). If the observed evolutionary change across replicates was dominated by parallel responses to shifting population densities, then the segregation of samples across the 2-D plane should, in part, correspond to the generation from which samples were derived. This analysis was conducted on mean, genome-wide F_st_ values, as well as F_st_ values computed separately for SNPs on each chromosomal arm. To quantify trends visualized using MDS, we translated the coordinates for each cage such that the centroid of all generation 0 samples was centered at the origin and then projected all points onto a single axis which was constructed by fitting a simple linear regression model to samples from generations 0-4 (regressions were constructed separately for each replicate cage). The resulting axis represents the primary axis of variation in the 2D plane during early expansion for each replicate (Fig. S5). We hypothesized that if sustained, directional selection imposed by population expansion was a primary driver of patterns in genomic variation, the samples from the remaining expansion generations would continue to proceed along the established axis of variation in the same direction. To evaluate the degree of concordance of early and late expansion generation samples we computed Pearson correlations between collection time and position along this axis. The significance of these correlations was determined via permutations of sample collection time point (*N =* 100 permutations). We conducted this analysis across all genome-wide SNPs, as well as separately for SNPs on each chromosomal arm. Finally, to disentangle whether differences in dynamics observed among chromosomal arms reflected true systematic differences in allele frequency behavior, as opposed technical artifacts driven by differences in the number of SNPs available for analysis across arms, we re-conducted this analysis using randomly sampled subsets of SNPs matching that present on each arm.

We quantified the extent to which individual SNPs exhibited parallel movement across cages and throughout expansion by fitting a generalized linear model to allele frequencies (formula: allele frequency ∼ expansion generation; *glm* function in base R) (34). Allele frequencies were weighted by the total number of chromosomes sequenced (N = 200) and depth per sample (60). This model assessed the significance of the linear relationship between allele frequency and generation of sampling across all cages, using a logistic link function and quasibinomial error model to reduce false positive associations (15,16,34). While an association between allele frequency and sampling timepoint within a single cage may represent either drift or selection, a significant association across all four replicate cages indicates allele frequency trajectories that are both predictable over time and parallel across populations. In this case, selection (or linked selection) is the more parsimonious explanation. P-values were adjusted using the Benjamini-Hochberg false discovery rate (FDR) correction (*p.adjust* package in base R). We considered a SNP significant if it exhibited an FDR < 0.01 and effect size > 2%. We further validated the parallelism inferred by the GLM by implementing a leave-one-out cross validation. Specifically, we iteratively identified sets of parallel SNPs (FDR < 0.01 and effect size > 2%) across expansion samples using a GLM and allele frequency data from three of the four replicate cages. We then quantified the frequency shifts of the favored allele at significant sites in the left-out cage between generation 0 and 9 of expansion. For sets of parallel SNPs identified via each iteration of the leave-one-out validation, we matched each parallel SNP to a control SNP based on chromosomal arm, starting frequency (within 5% of generation 0 frequency), inversion status, and recombination rate. We compared the distribution of allele frequency shifts at parallel vs. matched control sites (using a two-tailed, paired t-test) for each left out cage to infer if the magnitude of allele frequency change exceeded background allele frequency movement, and whether the dominant direction of allele frequency change was in a direction concordant with the three training cages. We conducted this analysis for SNPs identified genome-wide, as well as separately for each chromosomal arm.

Finally, we leveraged the genomic distribution of GLM significant SNPs to disentangle the minimum number of independently selected loci from the coordinated signal across SNPs expected as a result of linkage and genetic draft. Specifically, we implemented a method developed and described by (16). In brief, this method computes scores of GLM significance in sliding windows of 500 SNPs, using a 100 SNP step size. An empirical FDR for each window is generated by comparing observed scores to those generated for permuted windows. Windows with an empirical FDR < 0.05 are defined as significantly enriched and overlapping enriched windows are merged into a single, enriched window. A set of independently segregating loci is the determined by computing average SNP-pair linkage (using the squared correlation of founder genotypes) between all non-overlapping windows within 1 Mb. Windows with average SNP-pair linkage is > 0.03 are merged (this value was empirically determined via chromosome-wide patterns of linkage between SNPs (14, 16)), and this process continues iteratively until no windows with high linkage remain. We have previously shown that this method of identifying independent loci based on patterns of physical linkage indeed produces sets of loci with independent allele frequency movement throughout evolution (14). Source code for this analysis is provided at https://github.com/greensii/dros-adaptive-tracking). We used the most significant SNP within each unlinked locus to compute selection acting across reproduction selection. Specifically, we used the observed allele frequency change to quantify selection coefficients as: s = Δp/(p x (1-p)) where p corresponds to the allele frequency of the focal marker SNP. We note this derivation assumes no impact of genetic drift on allele frequencies and a constant selection coefficient throughout expansion, reasonable assumptions in the context of the large populations and short timescale over which we conducted the experiment and the controlled, isolated selection on reproductive output throughout this period.

### Quantifying evidence of fecundity and desiccation tolerance trade-offs

We next used samples collected throughout truncation, during which patterns of survival and selection were most likely underpinned by differences in desiccation resistance among genotypes, to quantify the presence of fitness-tradeoffs at those alleles systematically favored during expansion. First, we re-conducted our MDS analysis of pairwise divergence values, this time including all samples collected across both phases of the experiment. While general qualitative inference may be obtained from the visualization of this analysis, we explicitly quantified the relative movement of expansion and truncation samples across this 2D plane to provide a more rigorous investigation into the relative direction of allele frequency movement across phases. Specifically, as above, we translated MDS coordinates for each cage such that the centroid of all hour 0 truncation samples was centered at the origin, and then projected onto a single axis which, as above, was constructed by fitting a simple linear regression model to samples from expansion generations 0-4. We hypothesized that if our contrasting reproduction/stress-tolerance selection environments were inducing genome-wide reversions in allele frequency movement, then the projection of samples collected throughout truncation would regress in the opposite direction as that observed for the expansion samples. The degree of concordance/discordance for truncation samples were quantified as the Pearson Correlation between collection time and position along this axis, and the significance of these correlations was determined via permutations of sample collection time point (*N =* 100 permutations). This analysis was also conducted both genome-wide and separately for each chromosomal arm and, as above, we used genome-wide subsampling to infer whether variation in dynamics among chromosomal arms was a technical artifact.

Finally, we aimed to corroborate patterns observed in the dimensionality reduction of genome and chromosome-wide F_ST_ values described above by accruing additional evidence of genome-wide trade-offs using allele frequency trajectories. We first isolated those SNPs with strong evidence of linkage to a selected locus throughout expansion (GLM FDR < 0.01 and allele frequency shift > 2%). We then quantified the behavior of the rising allele at each SNP throughout the progression of truncation, and evaluated whether these alleles were more likely to exhibit a reversion in trajectory direction, or continue in the same direction, using a Chi-squared test and set of control SNPs (matched on chromosome, starting frequency, recombination rate, and inversion status). We further assessed if and how the results of this analysis varied when only evaluating trajectories of expansion-identified SNPs that also exhibited independent evidence of linkage to a selected locus (GLM FDR < 0.05 and allele frequency shift > 1%) during truncation. Finally, we further quantified the distribution of expansion-identified SNPs, genome-wide and per chromosomal arm, using empirical cumulative distribution functions (eCDF). The eCDFs for SNPs with sustained directional movement across treatments and evidence of fluctuating selection were evaluated independently via comparison to eCDFs generated via a set of matched control SNPs, and using a Kolmorgorov-Smirnov test (*stats* package in R). We then used the set of 250 unlinked loci identified throughout to further quantify the signal of fluctuating selection in our data.

Specifically, for each locus, we computed the mean frequency shift of expansion favored alleles (GLM FDR < 0.05 effect size > 2%) during truncation. We then generated a set of allele frequency shifts using matched control SNPs for each locus (matched on starting frequency, chromosome, inversion status, and recombination rate) and a paired, two-sided t-test to infer whether the locus continued in a consistent direction during truncation (positive median shift; t-test FDR < 0.05), was neutral, or moved significantly in the opposite direction (negative median shift; t-test FDR < 0.05).

### Assessing the role of inversions in fitness trade-offs

From the known cosmopolitan inversions found in the DGRP lines (62), we analyzed those that occurred in at least 10 of our founding strains (i.e., > 4%). We used chromosome-level haplotype frequencies to compute the frequencies of each of these inversions independently during expansion and truncation. Next, we regressed inversion frequencies through time using a generalized linear model (logistic link function, and quasibinomial error variance) to classify inversions moving systematically across replicates throughout each phase (Benjamini-Hochberg corrected P-value < 0.1). For those inversions displaying significant frequency change across both expansion and truncation, we characterized those that exhibited thus fluctuating selection and trade-offs (i.e. switched directions) across phases.

We next aimed to quantify the extent to which SNPs within, or in tight linkage of, of the assayed inversions underpinned genome-wide signals of trade-offs quantified in our experiment. We thus first generated a winnowed set of genome-wide SNPs, eliminating those either within, or up to 100 Kb away from, inversion breakpoints. We then explored whether we retained signals of parallel adaptation throughout expansion and trade-offs during truncaiton using this inversion-free SNP panel and re-conducting our F_ST_ -based MDS analysis of expansion and truncation samples (described in *Quantifying evidence of fecundity and desiccation tolerance trade-offs*, above).

### Quantifying the emergence of trade-offs during adaptation to natural environmental fluctuations

We next quantified whether alleles identified via selection under sustained population expansion in a laboratory setting can predict patterns of adaption and trade-offs in an outdoor environment. Specifically, we leveraged data from an independent study year when we monitored patterns of genomic variation in a genetically diverse, outbred population that was split into a series of large, replicate cages maintained in a controlled, indoor laboratory, as well as outdoor mesocosms exposed to natural environmental fluctuations. Allele frequency data for the outdoor mesocosms were previously described, analyzed, and reported in Bitter *et al.* (2024). Briefly, the outdoor mesocosms were located in Philadelphia, Pennsylvania and consisted of twelve, replicate 2 m^3^ cages, each consisting of a single dwarf peach tree and exposed to natural environmental fluctuations. The paired, indoor cage study (not reported or analyzed by Bitter *et al.* (2024)) consisted of ten, replicate ∼0.5 m^3^ cages housed in a temperature-controlled laboratory at the University of Pennsylvania.

The replicate cages for both the outdoor mesocosms and indoor cage study were seeded with a genetically diverse, outbred population, derived from a panel of 76 inbred strains originally collected wild from Linvilla Orchards, Media, PA (Supplementary Data File 2).

Constat food was supplied to the replicates within each environment, whereby four hundred ml of Drosophila media (‘Spradling cornmeal recipe’) was provided in 900 cm^3^ aluminum loaf pans within each cage three times per week. These pans provided the only source of food and egg laying substrate, and egg laying upon each loaf pan was carried out for two days, after which a new pan was added, and the original pan was covered with a mesh lid to prevent any further laying. The lids were removed after eclosure was first observed, causing each replicate to experience a near continual input of new flies that evolved with overlapping generations. The census size of the adult population in the outdoor mesocosms were estimated five times throughout the progression of the experiment (19 July, 5 August, 20 August, 17 September, and 20 October) (14), while the final census in the indoor cages were estimated at the end of the monitoring period via collection and volumetric quantification of all flies remaining in each replicate. The outdoor mesocosms were sampled to quantify patterns of genomic variation weekly for 9 weeks (13 July – 7 September), after which samples were collected on 21 September, 20 October, and following the first freeze and population crash on 20 December (12 total time points). The indoor cages were sampled at the first, second, eighth, and eleventh time points. The overlapping monitoring period of the outdoor and indoor cages encompassed the rapid population expansion and ultimate stabilization of cage densities as the replicates in each environment presumably reached carrying capacity (Fig S25) (14). The selective landscape experienced during this shared monitoring period probably mirrors that experienced under the sustained population expansion imposed by population expansion portion of the first experiment of this study, described above. The notable differences between these experiments, however, is that the first experiment increased food availability as a function of population density throughout expansion, while our latter experiment maintained a constant food substrate throughout the study period. Thus, this paired, indoor-outdoor study likely also imposed selection on traits impacted by density regulation (e.g., larval competition). Analogous to the truncation portion of the first experiment, the final collection interval of the outdoor mesocosm study (time point 11->12) encompassed the onset of winter, during which time a total crash of the adult populations in the cages was observed. Selection during this phase of the experiment was thus hypothesized to favor increased in stress tolerance traits associated with the deteriorating abiotic conditions.

Methods for sampling flies for sequencing from the outdoor mesocosms and indoor cages were similar to those described above: eggs were collected overnight directly from each replicate cage, and larval development was carried out in the indoor environment where eggs developed and eclosed to F1 adults in 30 cm^3^ cages. A random set of 100 females were sampled from each cage 3-5 days post-eclosure and preserved in 99% ethanol at -20° C (14). Genomic DNA was extracted from pools of flies for each replicate and time point using the Monarch Genomic DNA Purification Kit (New England Biolabs). Libraries were nonstructured using the Illumina DNA Prep Tagmentation Kit and all samples were sequenced on Illumina Novaseq 6000 flow cells using 150 bp, paired end reads. Raw sequencing reads from each sample were trimmed of adapter sequences and bases with quality score < 20, and aligned to the *Drosophila melanogaster* v5.39 reference genome using bwa and default parameters(58). Aligned reads were deduplicated using Picard tools (http://broadinstitute.github.io/picard/) and the final set of reads for each sample was down-sampled to obtain an equivalent, genome-wide coverage of 8x across all samples (>100x effective coverage) (14,60). Haplotype informed allele frequencies from reads from each replicate and time point were then generated using the local inference pipeline described above (see *Allele Frequency Calculation*) (59,60). The final set of allele frequencies only included those sites with an average minor allele frequency > 0.02 in the baseline population, and present in at least one evolved sample at a MAF > 0.01, ultimately yielding 1.9 M SNPs (14).

We quantified the extent to which evolution in the outdoor environment is driven by the selective pressures solely associated with increasing population density (e.g., selection for increased fecundity, faster developmental rate, increased competitive ability). Specifically, we identified alleles with systematic behavior independently in the indoor and outdoor cages using GLM (FDR < 0.01; allele frequency change > 2%), and compared the observed overlap between these lists using a Chi squared test and with null expectations generated via a set of matched control SNP set. Next, we queried whether the dominant direction of selection throughout the shared monitoring period of the indoor and outdoor environment was consistent. Specifically, we quantified the mean frequency shift of the rising allele at SNPs identified in the indoor environment via GLM, in the outdoor cages during the same time interval. We determined whether the magnitude of frequency shifts exceeded background allele frequency movement expected via matched control SNPs, and whether the dominant direction was parallel or anti- parallel given the sign of observed median shift (analysis conducted both genome-wide and separately for each chromosomal arm using a paired t-test and sign test). Finally, to test whether the total population crash in the outdoor mesocosms elicited evidence of the trade-offs, we quantified the behavior of the indoor identified alleles throughout this period. Through this, we used a common set of alleles, determined as linked to an advantageous locus during increasing population density in a controlled, lab-based setting, to infer the presence of trade-offs during a bout of population expansion and collapse in a semi-natural setting.

## Acknowledgements

We are grateful to members of the Petrov and Schmidt labs for discussion during experimental design, lab work, and data analysis. We thank our funding organizations: the National Science Foundation (NSF PRFB 2109407 to M.C.B.) and the National Institutes of Health (NIH 5R35GM118165-07 to D.A.P and NIH R01GM100366 and R01GM137430 to P.S.), as well as funding support from the Chan Zukerberg Biohub.

## Financial Disclosures

National Science Foundation (NSF PRFB 2109407 to M.C.B.) and the National Institutes of Health (NIH 5R35GM118165-07 to D.A.P and NIH R01GM100366 and R01GM137430 to P.S.), as well as funding support from the Chan Zukerberg Biohub. The funders had no role in study design, data collection and analysis, decision to publish, or preparation of the manuscript.

## Statement of Authorship

The experiment was conceived by D.A.P, P.S, A.O.B., S.R., and M.C.B. Data curation for the population expansion/truncation experimental evolution study was conducted by S.R., N.B., S.T., A.O.B., S.G., and P.S. Data curation paired indoor and outdoor mesocosm study was conducted by M.C.B., S.B., H.O., and P.S. Formal analysis was carried out by S.G., J.H., E.L., and M.C.B. The original manuscript was prepared by M.C.B., S.G., P.S., and D.A.P. All authors reviewed and edited the final version of the manuscript.

## Data and code availability

Sequencing data from the DGRP lines used in the population expansion/truncation experiment are publicly available at http://dgrp2.gnets.ncsu.edu/data.html. Haplotype-informed allele frequency estimates from evolved, outbred samples and used in statistical analysis will be available at upon article publication. Founder line sequences for the paired indoor/outdoor mesocosm study are available at NCBI Accession PRJNA722305. Outdoor mesocosm sequences generated via pooled sequencing for this experiment are available at NCBI accession PRJNA1031645 and allele frequency data is available at the following Dryad repository: https://doi.org/10.5061/dryad.xd2547dpv. Indoor sequences and allele frequency data will be available upon article publication. Code associated with all analyses conducted in this manuscript are publicly available at the following repository: https://github.com/MarkCBitter/Drosophila-fitness-trade-offs.

## Supplementary Information for Bitter *et al.* (2024)

### Supplementary Methods

To evaluate whether patterns of linkage between loci, antagonistic pleiotropy at a single SNP, and/or haplotype-level antagonistic pleiotropy could generate the patterns of allele frequency change observed in our data we ran Wright-Fisher simulations with SLiM (1). First, we simulated the behavior of a single SNP with antagonistically pleiotropic behavior evolving under fluctuating selection. Specifically, in a population of size N=1000 diploid individuals, we introduced a target SNP ‘A’ at a range of initial, intermediate starting frequencies (p_0_ ∈ 33-64%). We traced the frequency of the favored allele over 13 generations: the first 9 generations corresponded to the expansion phase of our experimental evolution, and the fitness of A during expansion was labeled *s_expansion_*. The last 4 generations corresponded to the truncation phase, with the fitness of ‘A’ labeled *s_truncation_.* In our simulations, we considered every possible combination of fitness values (*s_expansion_*, *s_truncation_*), where *s_expansion_* was varied between 1.0 (neutral) and 1.2 (highly advantageous), and *s_truncation_* was varied between 1.0 (neutral) and 0.8 (highly deleterious). This range of selection coefficients was based on the empirical selection coefficient values we computed from the allele frequency data in our reproduction/stress-tolerance selection experiment (**Fig. S8**). In all simulations, here and below, we assumed no dominance (h=0.5). In total, we ran 250 replicate simulations for each combination of two selection parameters (s_expansion_, s_truncation_).

We then ran a set of simulations in which two separate SNPs, ‘A’ and ‘B’, are under selection during expansion or truncation, respectively. The two SNPs are located a set distance, *d*, nucleotides apart. We seeded the genotypes of individuals in the initial generation by fixing initial frequencies of alleles, *p*_0_^(A)^ and *p*_0_^(B)^, as well as their linkage state. The linkage was determined by the magnitude of the LD measure r^2^ between A and B, with an r^2^ < corresponding to an effectively "unlinked" state between A and B, and r^2^>0.05 corresponding to A and B being "highly linked”. We further distinguished between the two different ways in which A and B could be highly linked by using the sign of r: the "raw" correlation between ‘A’ and ‘B’ that, when squared, yields *r*^2^. When *r* > 0 and *r*^2^ > 0.05, we consider A and B to be both highly linked and in "attraction", meaning that favored alleles (those that are assigned fitness effects in the simulations) at both loci are assorted together.

Conversely, when *r* < 0 and *r*^2^ > 0.05, ‘A’ and ‘B’ are highly linked and in a state of "repulsion", whereby the favored allele of ‘A’ is assorted together with the disadvantageous allele of ‘B’, and vice versa. To summarize, we have distinguished three linkage states: "unlinked", "highly linked in attraction", and "highly linked in repulsion".

As the range of possible values of *r* and *r*^2^ depends on the initial frequencies of A and B (2), for each simulation we sampled the value of *r* uniformly from its range of possible values.

For example, when simulating loci in attraction, we chose *r* uniformly in the interval (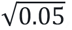, *r_max_* (*p*_0_^(A)^, *p*_0_^(B)^)). When simulating loci in repulsion, we chose *r* uniformly in the interval (*r_min_* (*p*_0_^(A)^, *p*_0_^(B)^), 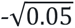). Finally, for unlinked loci, we chose *r* randomly in the interval (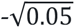, 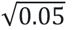). The functions *r_min_* and *r_max_* were derived following the approach of VanLiere and Rosenberg (2008). After *r* was chosen, we computed the numbers of individuals possessing haplotypes AB as:

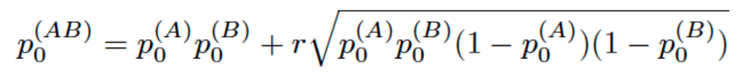

and used this value in setting the initial generation of the simulation.

After seeding the simulation by fixing initial frequencies of A and B and their linkage state in the first generation, we ran the simulation for 13 generations as above, with the first 9 generations corresponding to "expansion" and then 4 generations of "truncation". In all simulations we set *s_expansion_*^(B)^= *s_truncation_^(^*^A)^ = 0. The recombination rate ρ was equal to ρ=2.39×10−8 per generation per base pair during expansion, and to ρ=0 during truncation. The initial frequencies, *p*_0_^(A)^ and *p*_0_^(B)^, were chosen randomly in each simulation from the interval (0.34, 0.66). For the rest of the parameters, we formed a parameter grid and considered every possible combination of parameter values (linkage state, *d*, *s_exp_*^(A)^, *s_trunc_^(^*^B)^) where d ∈ {0.5, 1, 2, 4, 20} measured in megabases, s_exp_^(A)^ ∈ {1.0, 1.05, 1.1, 1.15, 1.2}, s_trunc_^(B)^ ∈ {1.0, 0.95, 0.9, 0.85, 0.8}, and linkage state was one of “unlinked”, “highly linked in attraction”, and “highly linked in repulsion”. We ran 1000 replicate simulations for each combination of parameter values. The seeding procedure described above ensures that we are quantifying average behavior of alleles in each linkage category with varying values of r^2^ and initial frequency. To summarize the results of the simulation, we computed maximum-likelihood estimates of “empirically observed” fitness values for A and B directly from the allele frequency changes (eq. (4) in (3)). For the expansion stage, we used all generations, while fitness during truncation was estimated only from the first generation. The estimate of fitness during the first generation of truncation represents the stress tolerance selection period of our experiment and explicitly quantifies the linkage effect of the B SNP on the A SNP’s allele frequency trajectory during this period.

Finally, to assess the net effect of linkage on the behavior of the allele frequency of SNP ‘A’ (that favored during expansion) during the first generation of truncation, we combined replicate simulation runs for all three linkage categories, and computed fitness of SNP A during the first generation of truncation from all combined simulations in which the attraction and repulsion states were equally likely to be observed. The resulting ‘net’ effect of linkage states on trajectories observed via this scenario then represents independent assortment of the ‘A’ and ‘B’ alleles.

### Supplementary Figures

**Figure S1.**
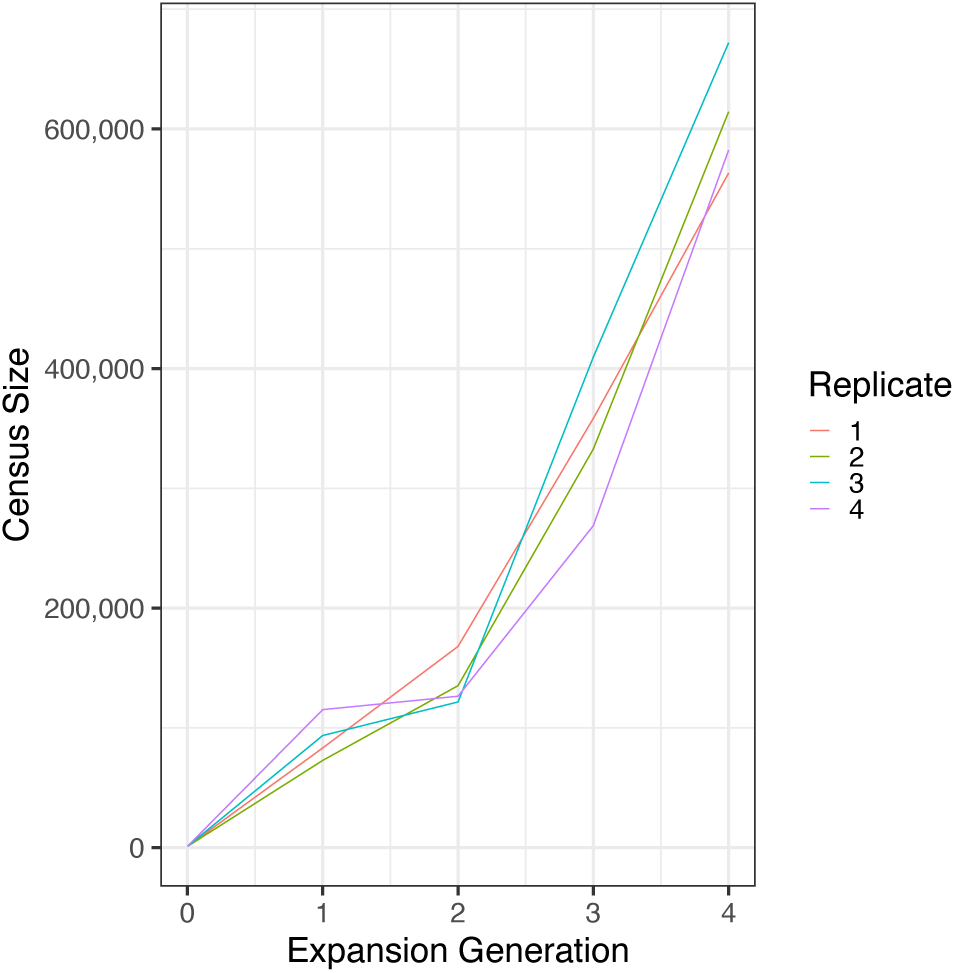
Census estimates of indoor replicate cages during the first four generations of population expansion.

**Figure S2.**
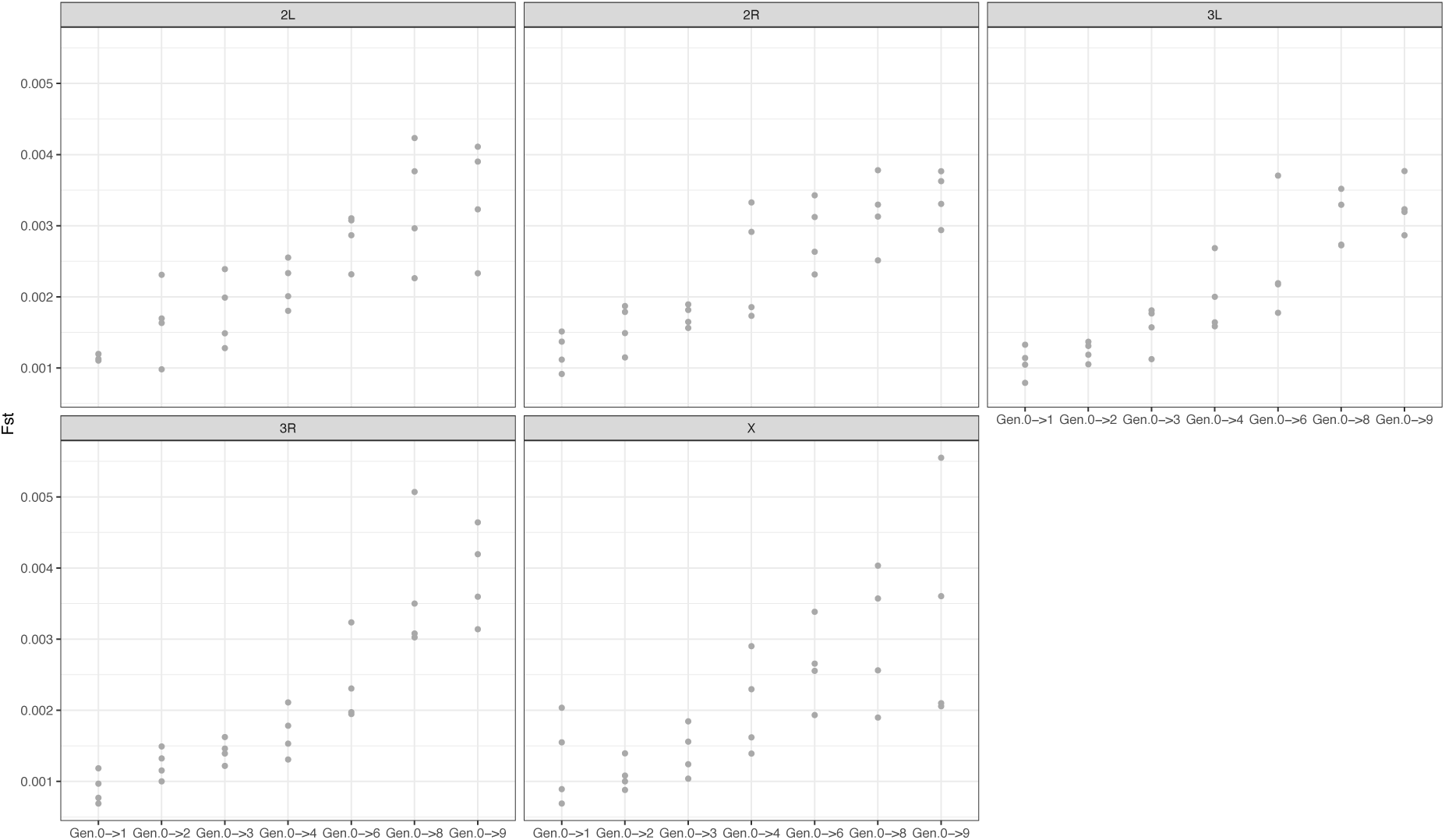
Evolution of allele frequencies across expansion generations. (A) Mean, genome-wide Fst between each evolved replicate and its generation 0 sample (Gen.0->n), segregated by chromosomal arm. Each grey dot corresponds to an individual replicate.

**Figure S3.**
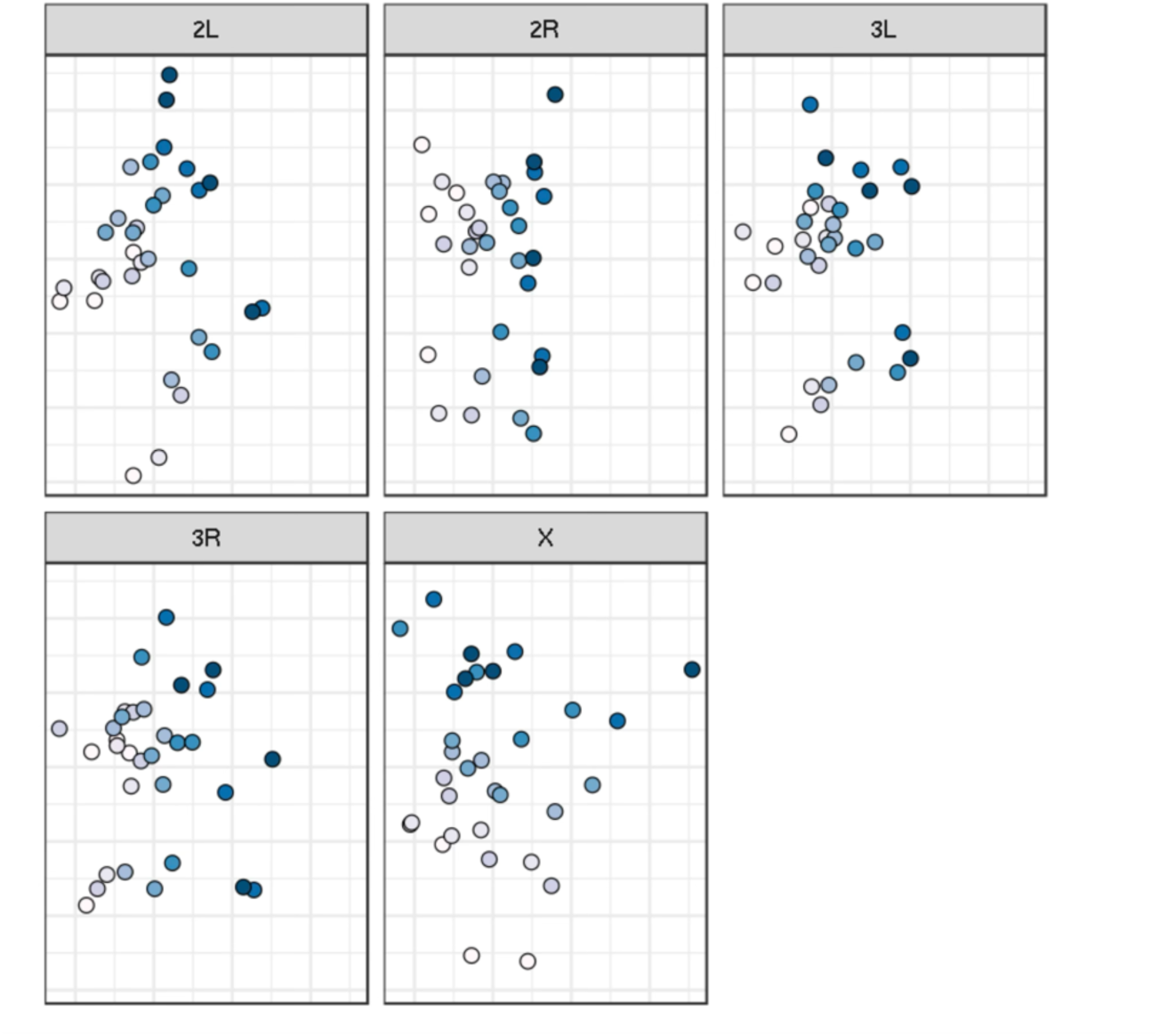
MDS of pairwise Fst values computed per chromosomal arm and across all expansion samples. Point Color corresponds to sample expansion collection generation.

**Figure S4.**
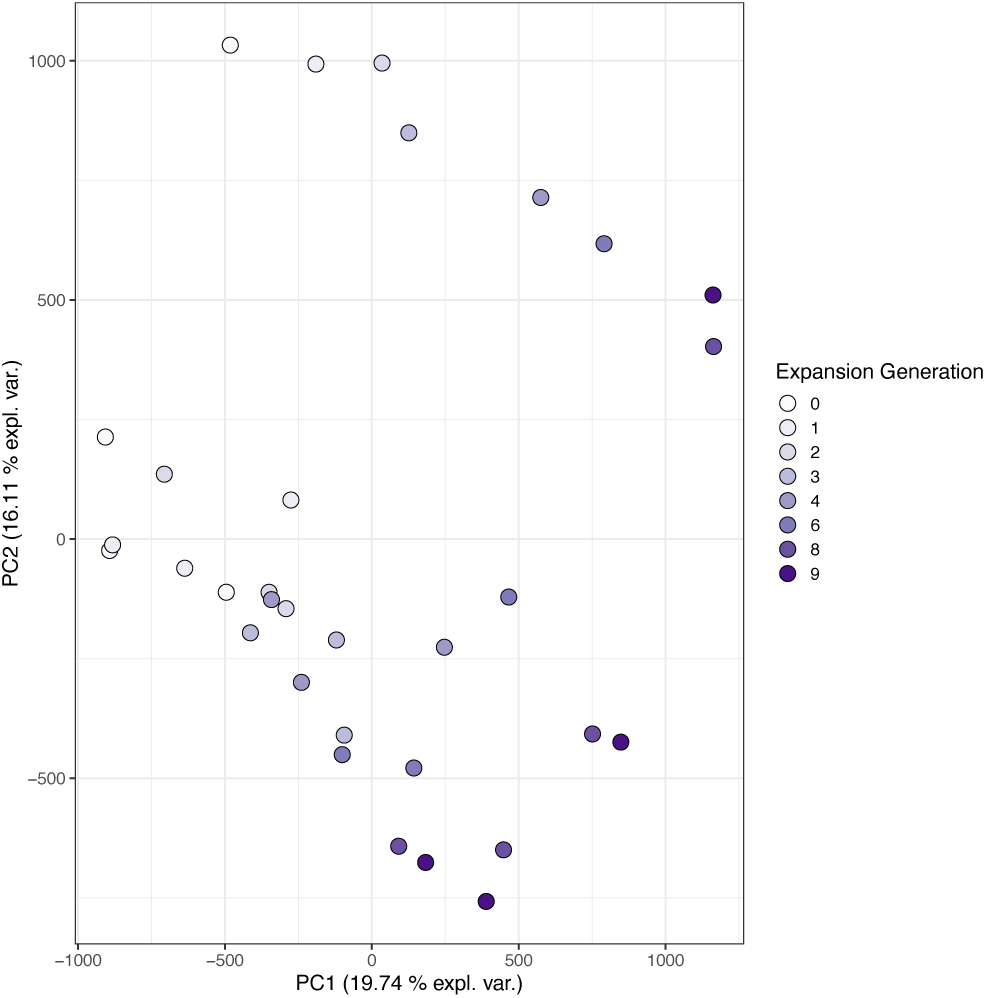
Principal component analysis of samples collected throughout expansion using allele frequency data across all 1.7 M SNPs. Samples are projected onto the first two principal components and colored in accordance with expansion generation collection time point.

**Figure S5.**
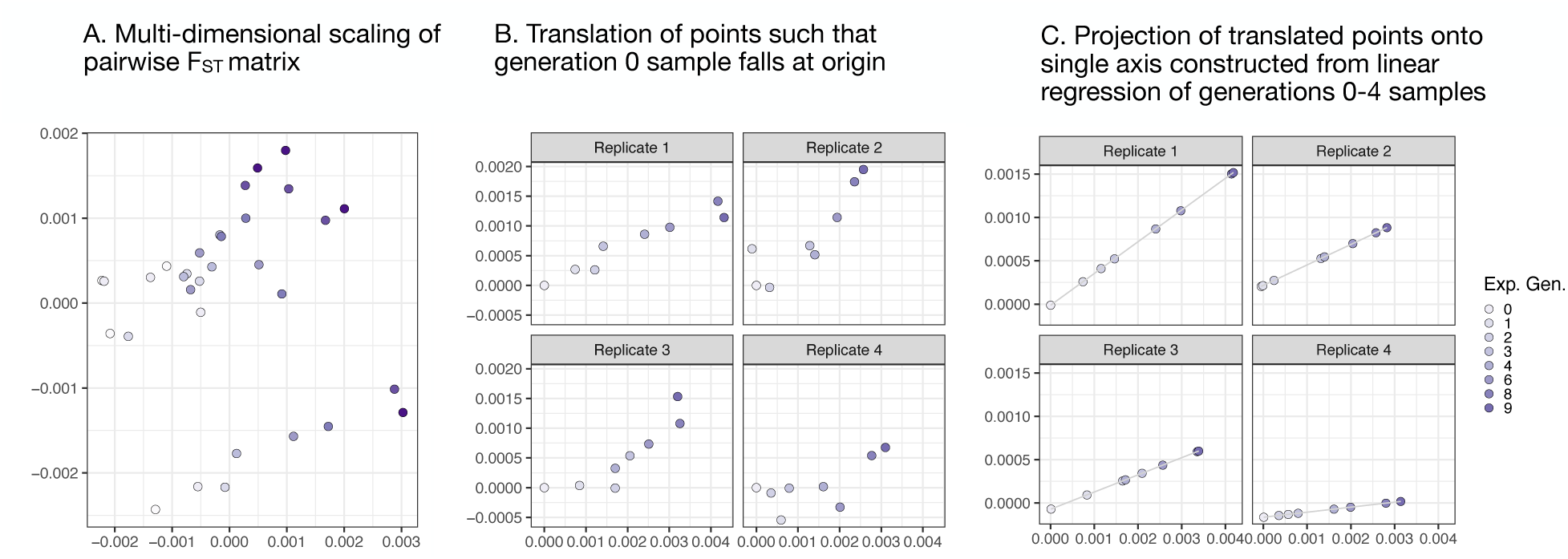
F_ST_-based MDS analysis workflow. (A) Multi-dimensional scaling of expansion sample pairwise F_ST_ matrix as in main figure 2. (B) Translation of expansion points such that, for each replicate, the generation 0 sample falls at the plot origin. (C) Projection of translated points (B) onto a single axis constructed via a simple linear regression of generation 0-4 translated points.

**Figure S6.**
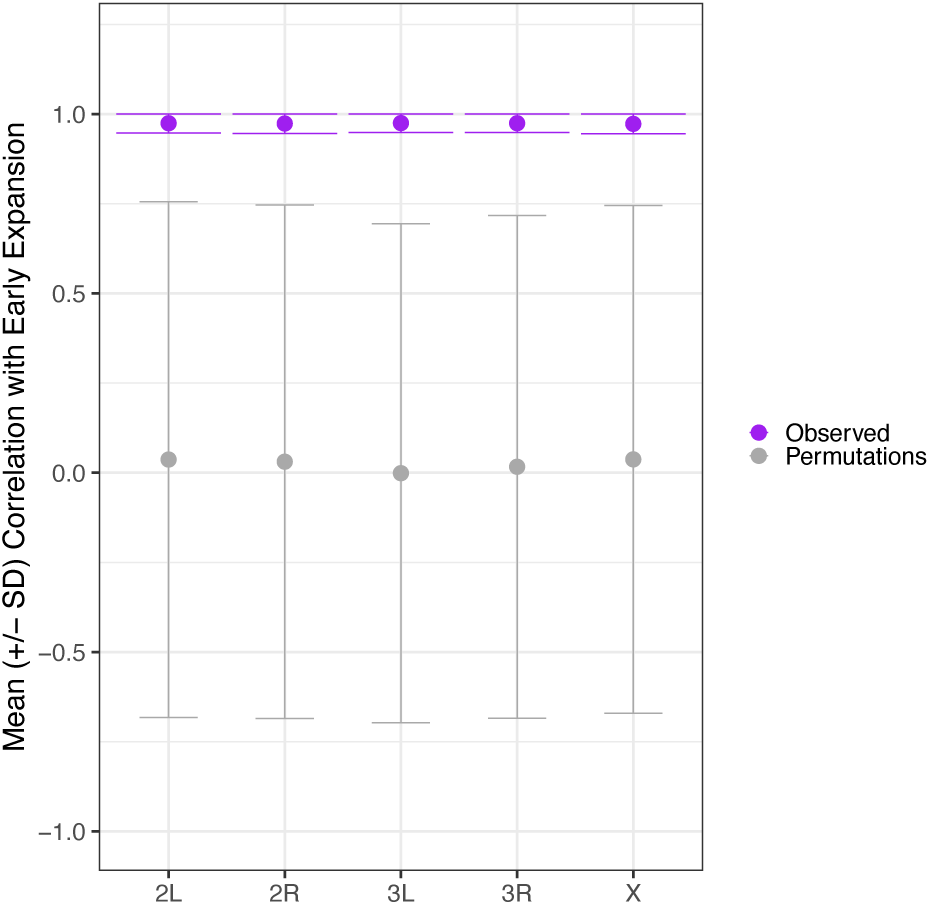
Average Pearson Correlation across replicates (+/- standard deviation) between sample expansion generation and distance along a one-dimensional axis constructed using F_ST_ MDS coordinates for early expansion (generation 0-4) samples. Purple points and error bars correspond to empirical values, while grey points and error bars correspond to values derived from empirical permutations (N = 100). Here, Fst values and MDS analysis was run on SNPs randomly sampled throughout the genome, matching the number of SNPs present on each chromosomal arm.

**Figure S7.**
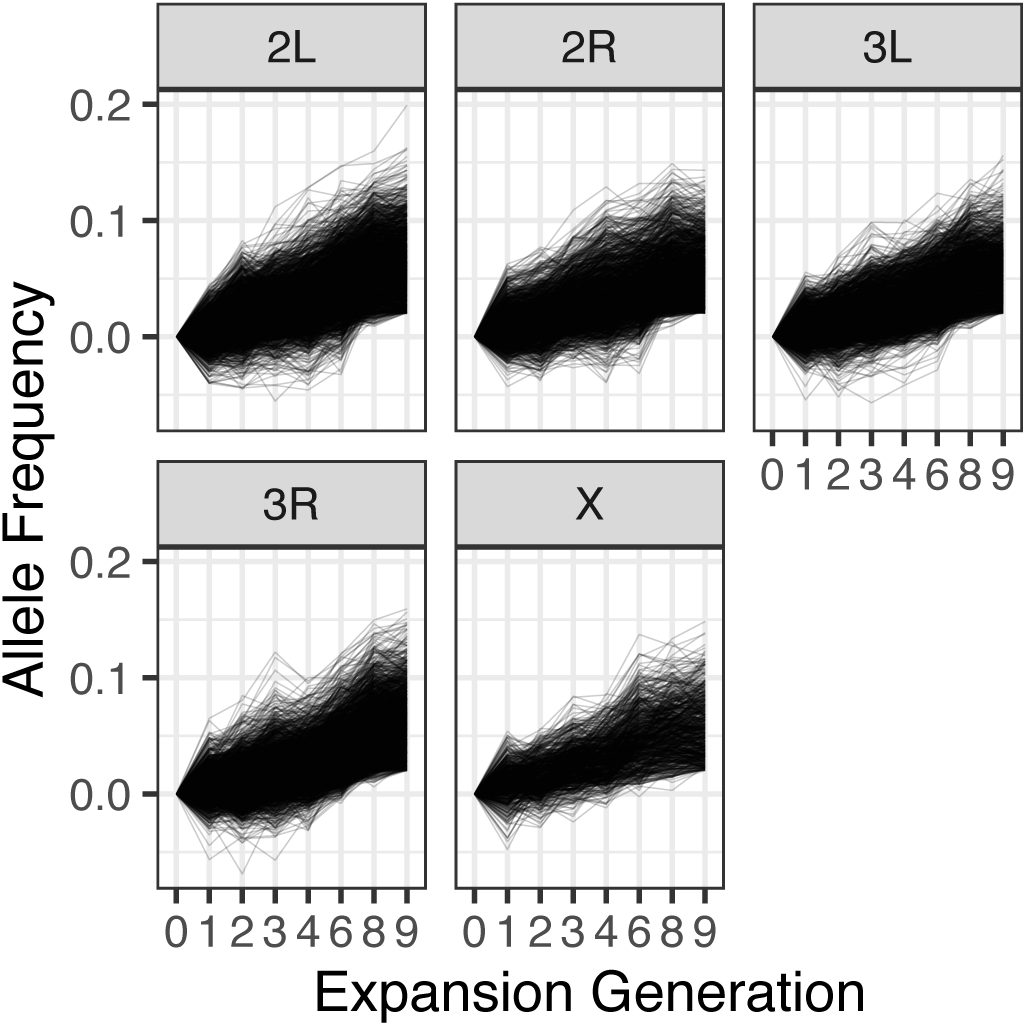
Trajectories of SNPs identified via GLM as exhibiting parallel frequency shifts across replicates throughout expansion (GLM FDR < 0.01 and effect size > 2%), segregated by chromosomal arm.

**Figure S8.**
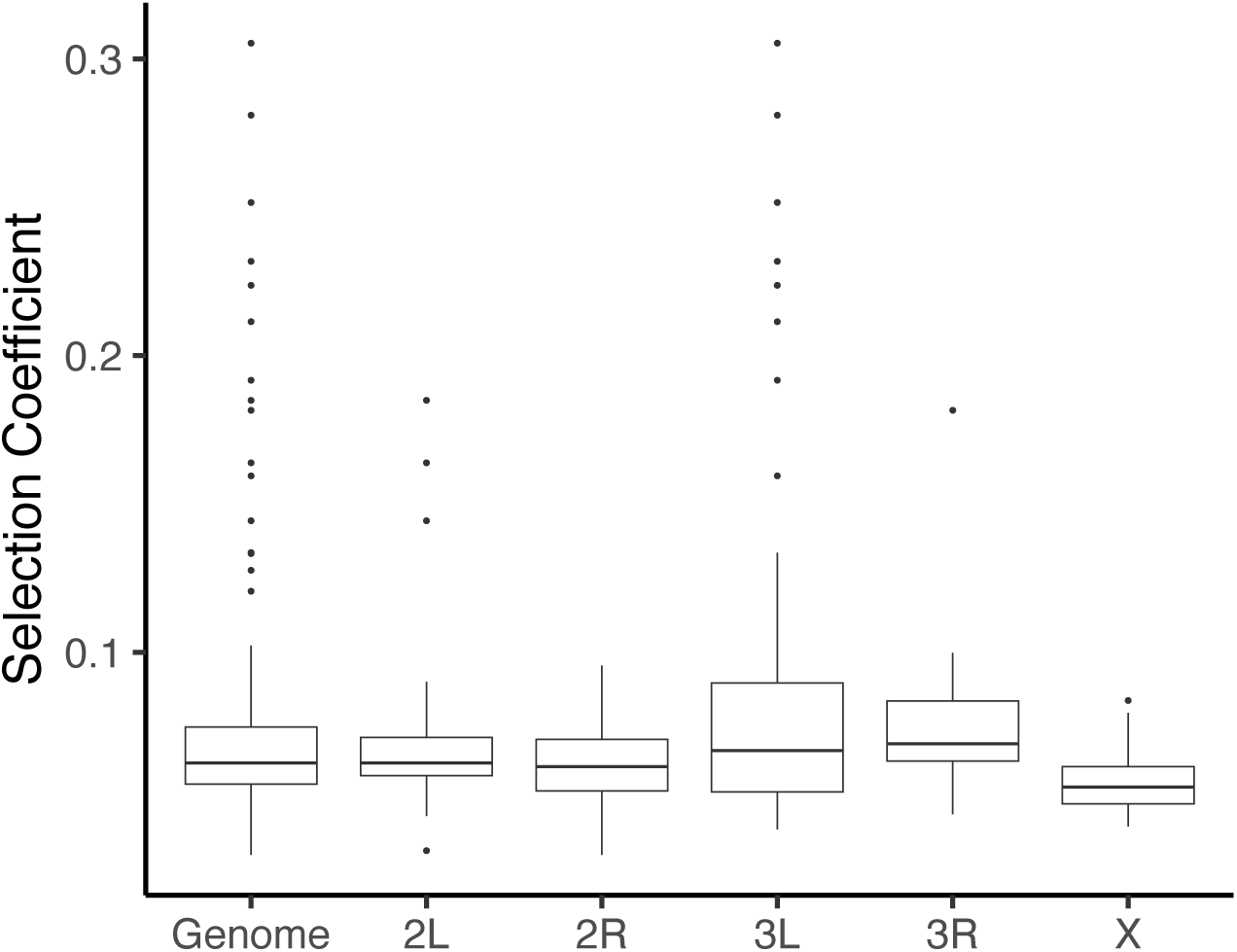
Per-generation selection coefficients computed using top marker SNP within each of the 250 unlinked loci identified throughout reproduction selection. Box plots span the 25th -75th percentile of distribution and horizontal bars indicate the median value (outliers depicted as dots).

**Figure S9.**
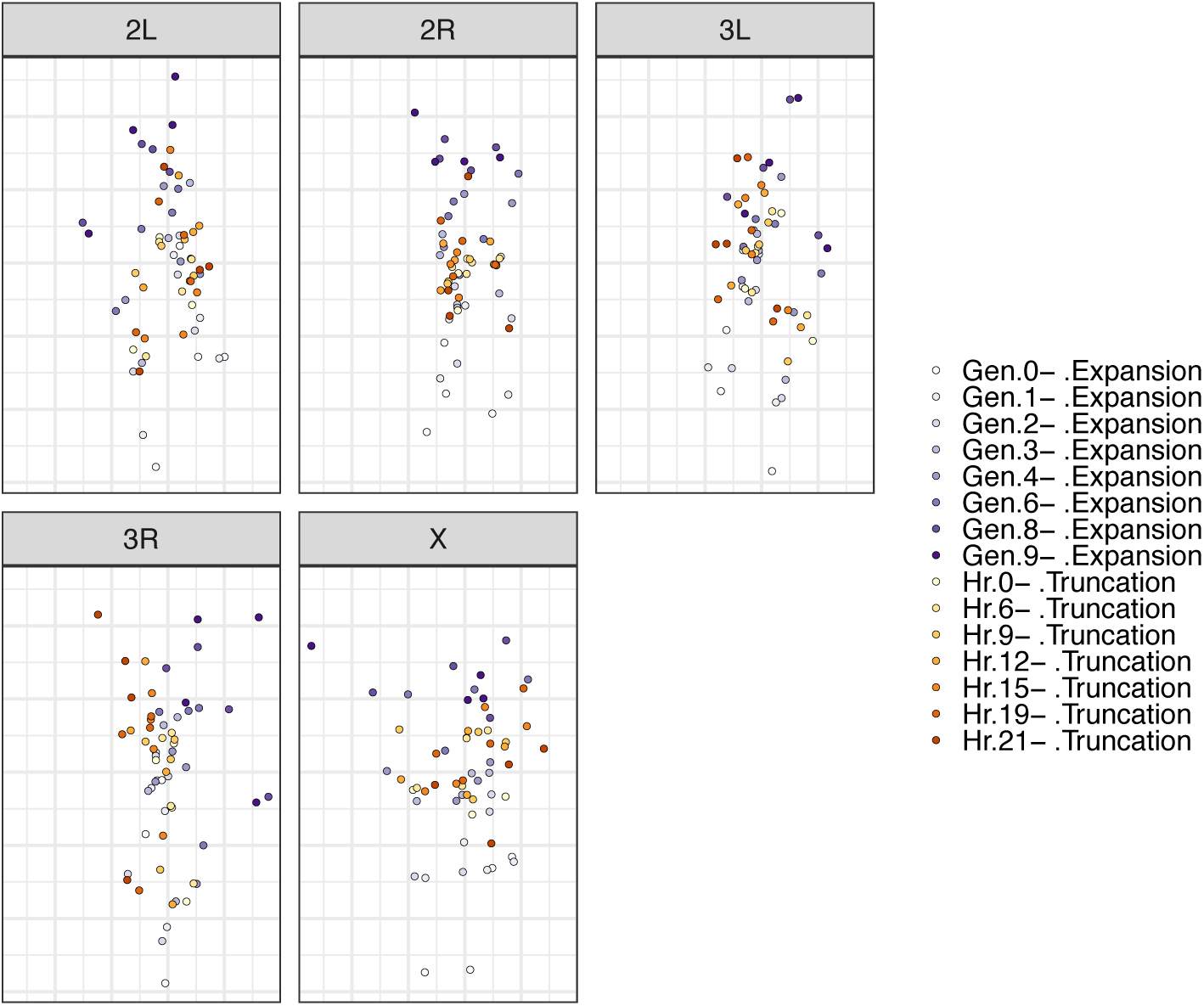
MDS of pairwise Fst values for each chromosomal arm and across all samples collected throughout expansion (purple-hue points) and truncation (orange-hue points). Points are shaded according to collection time point (darker hues indicate later expansion or truncation sampling generation/hour).

**Figure S10.**
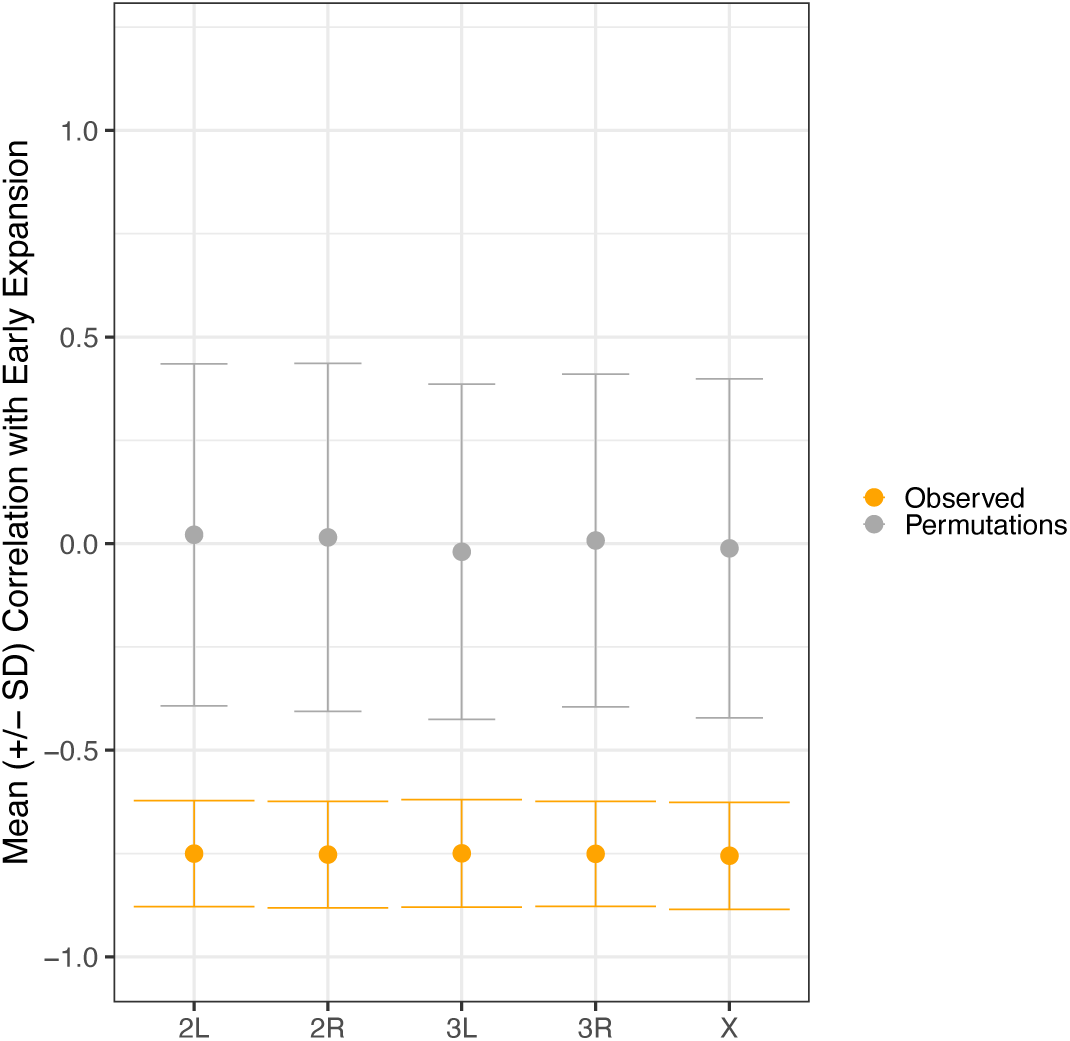
Average Pearson Correlation across replicates (+/- standard deviation) between sample truncation hour and distance along a one-dimensional axis constructed using F_ST_ MDS coordinates for early expansion (generation 0-4) samples. Orange points and error bars correspond to empirical values, while grey points and error bars correspond to values derived from empirical permutations (N = 100). Here, Fst values were derived from randomly sampled SNPs throughout the genome, matching the number of SNPs present on each chromosomal arm.

**Figure S11.**
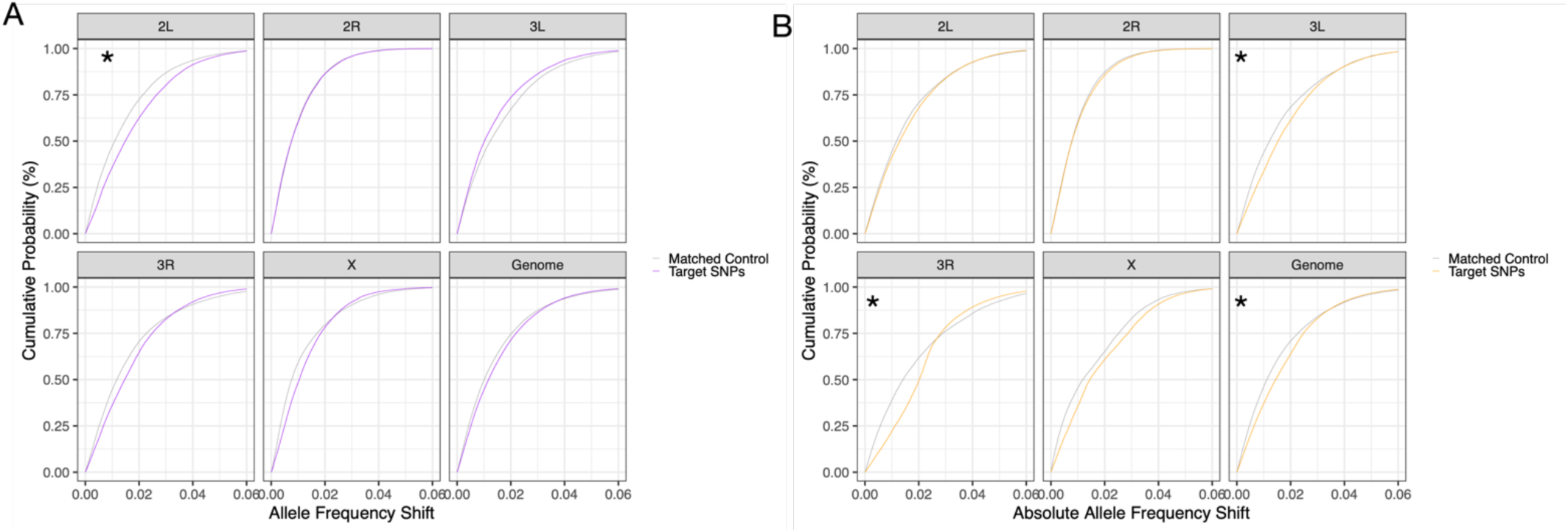
eCDFs of expansion-favored SNPs during truncation, segregated by those with sustained direction of movement (A) or reversals in direction (B). Asterisks correspond to instances in which the observed eCDF differed significantly from that generated via matched control SNPs (grey lines) (one-tailed Kolmogorov-Smirnov test, p-value < 0.05).

**Figure S12.**
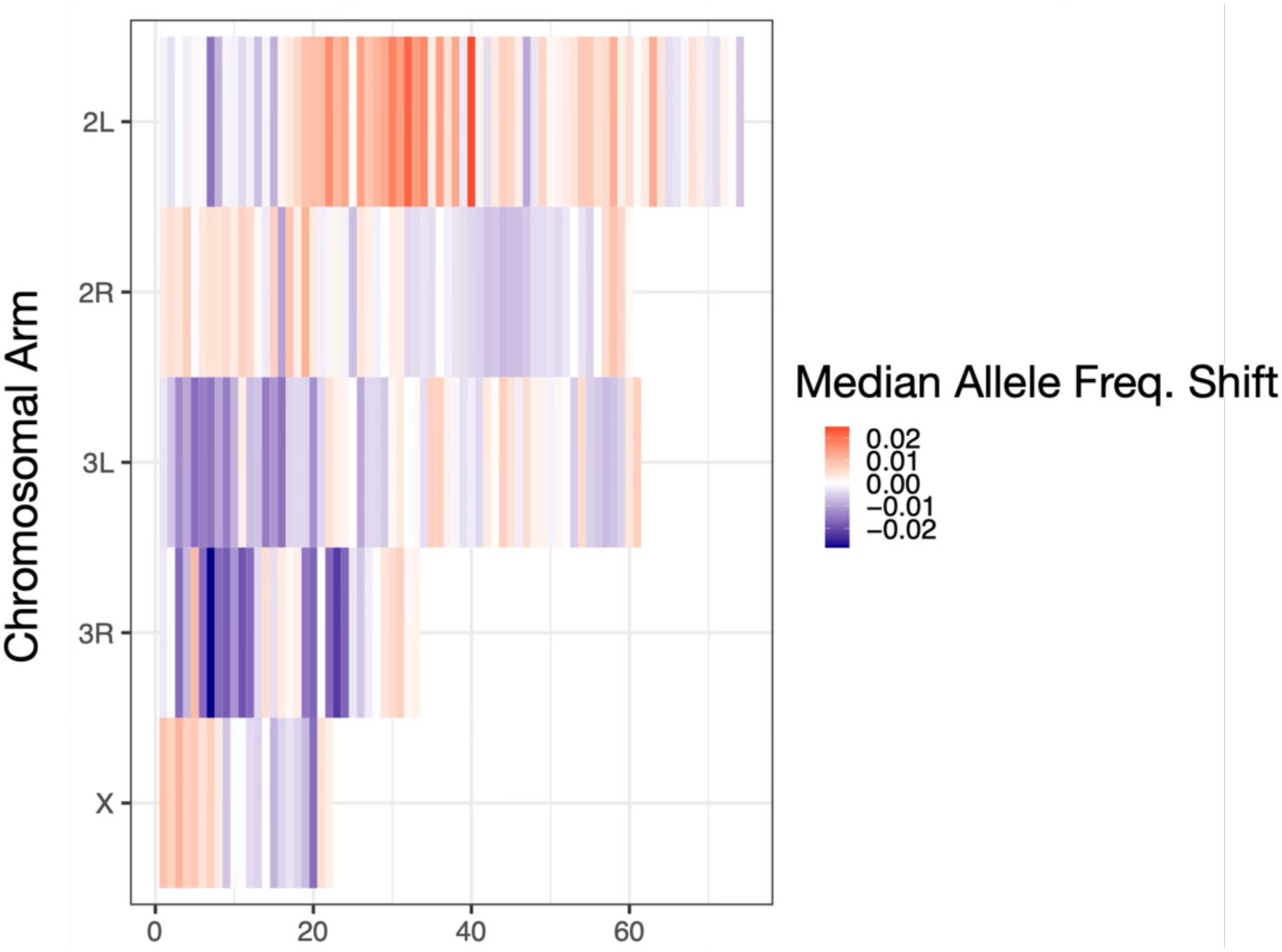
Expansion-favored locus trajectories (N = 250) throughout truncation. Loci are ordered according to chromosomal arm (y-axis) and position along chromosome (x-axis). X-axis ticks refer to the total number of loci identified on each arm. Tile colors correspond to the median allele frequency shift of each locus during survival stress tolerance selection. In total, 90 loci were inferred neutral during stress tolerance selection, while 90 exhibited signals of fluctuating selection (negative median allele frequency shift and paired T-test FDR < 0.05) and 70 under sustained directional selection during truncation (positive median allele frequency shift and paired T-test FDR < 0.05).

**Figure S13.**
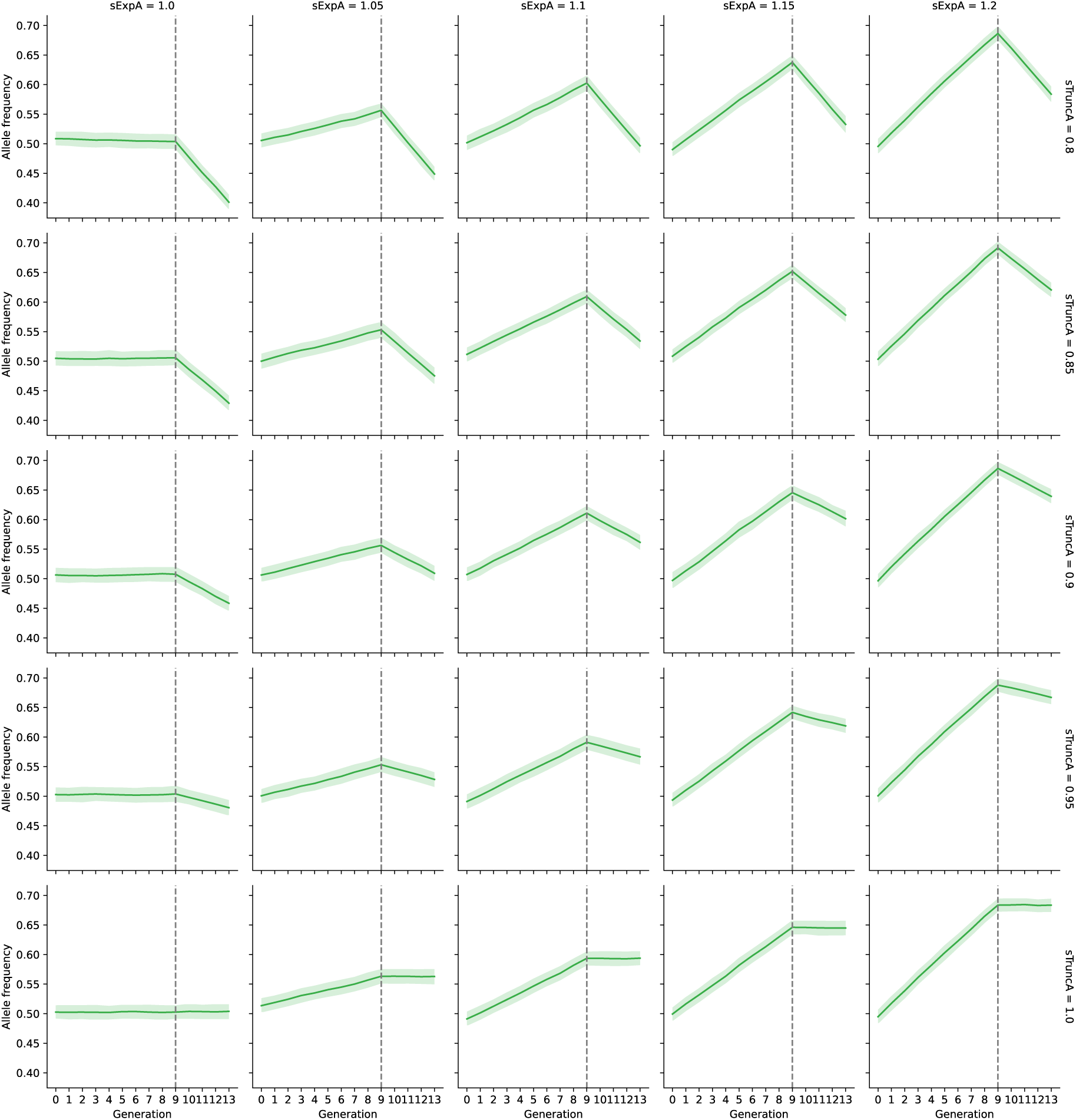
Simulated allele frequency trajectories of a SNP (‘A’) under a scenario of antagonistic pleiotropy across expansion (generation 0-9) and truncation (10–13). The selection coefficient of ‘A’ is ranged between 1.0 and 1.2 during expansion (columns) and (0.8 and 1.0) during truncation (rows). The solid line corresponds to the mean value across simulations and shaded area to the 95% confidence intervals (N = 250 total simulations per set of parameter values).

**Figure S14.**
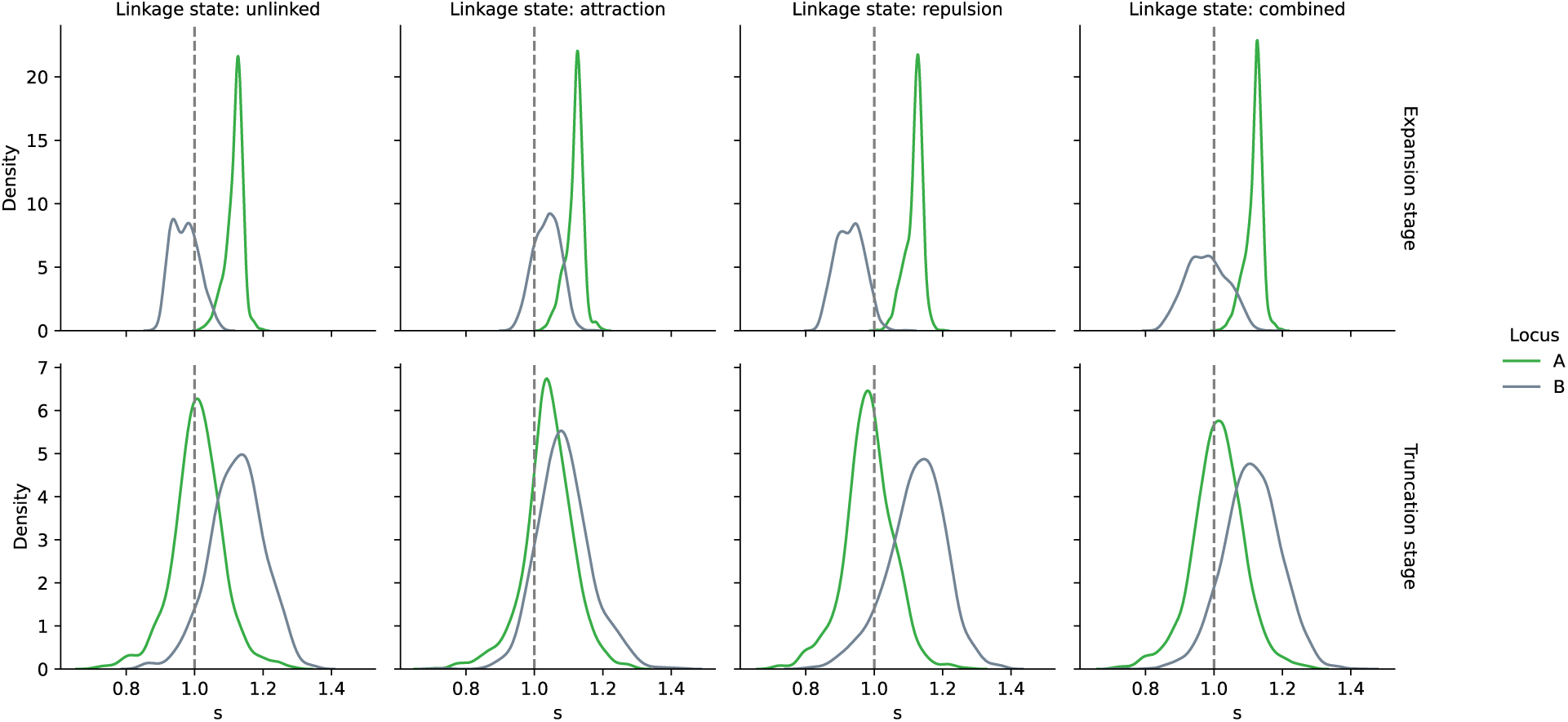
Kernal density estimates of empirical selection coefficients (s) computed during the first generation of truncation selection in two-locus SLiM simulations (N = 1000 simulations per scenario). In each case, the A and B loci are 1 Mb away, with locus A selected during expansion (fitness of favored allele during expansion = 1.1), but neutral during truncation, while B is neutral during expansion and favored throughout truncation (fitness of favored allele during truncation = 1.1). The columns represent different scenarios of linkage between the two loci, whereby ‘attraction’ represents a scenario in which the favored alleles at each locus are non-independently assorted with each other, and a state of ‘repulsion’ represents a scenario in which the favored allele at each locus was non-independently assorted with the disadvantageous allele at the other locus. The ‘combined’ scenario represents the average behavior across linkage scenarios.

**Figure S15.**
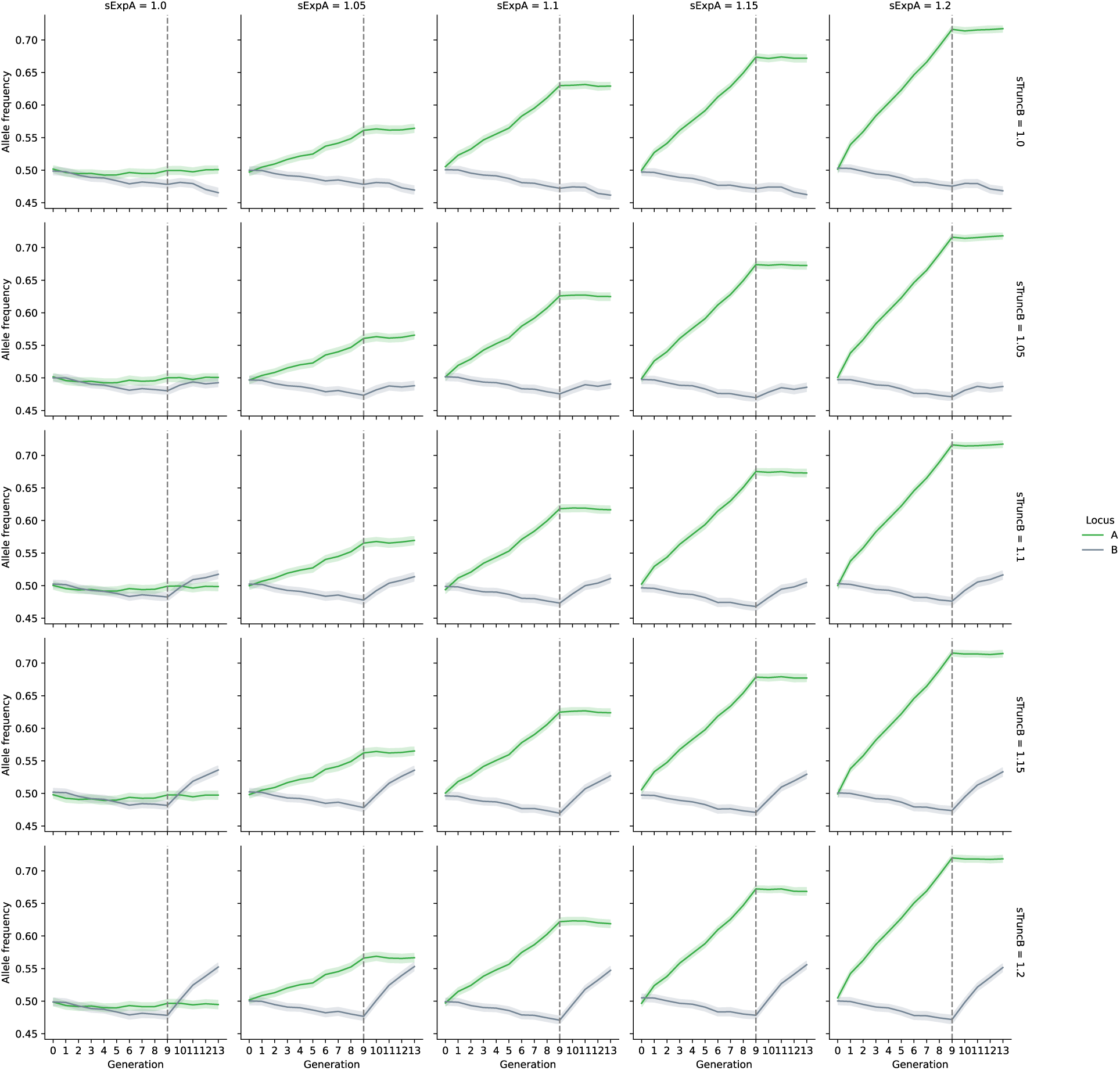
Simulated allele frequency trajectories for two unlinked SNPs under differential selection across expansion and truncation. The ‘A’ allele is under directional selection during the expansion period (generations 0-9) and neutral during the truncation phase (generations 10-13). The ‘B’ allele is neutral during expansion, then comes under directional selection during truncation. The selection coefficient of ‘A’ is ranged between 1.0 and 1.2 during expansion (columns) and remains 1.0 during truncation. The selection coefficient of ‘B’ is constant at 1.0 during expansion and ranged between 1.0 and 1.2 during truncation (rows). The solid line corresponds to the mean frequency value of each allele across simulations and shaded area to the 95% confidence intervals (N = 250 total simulations per set of parameter values).

**Figure S16.**
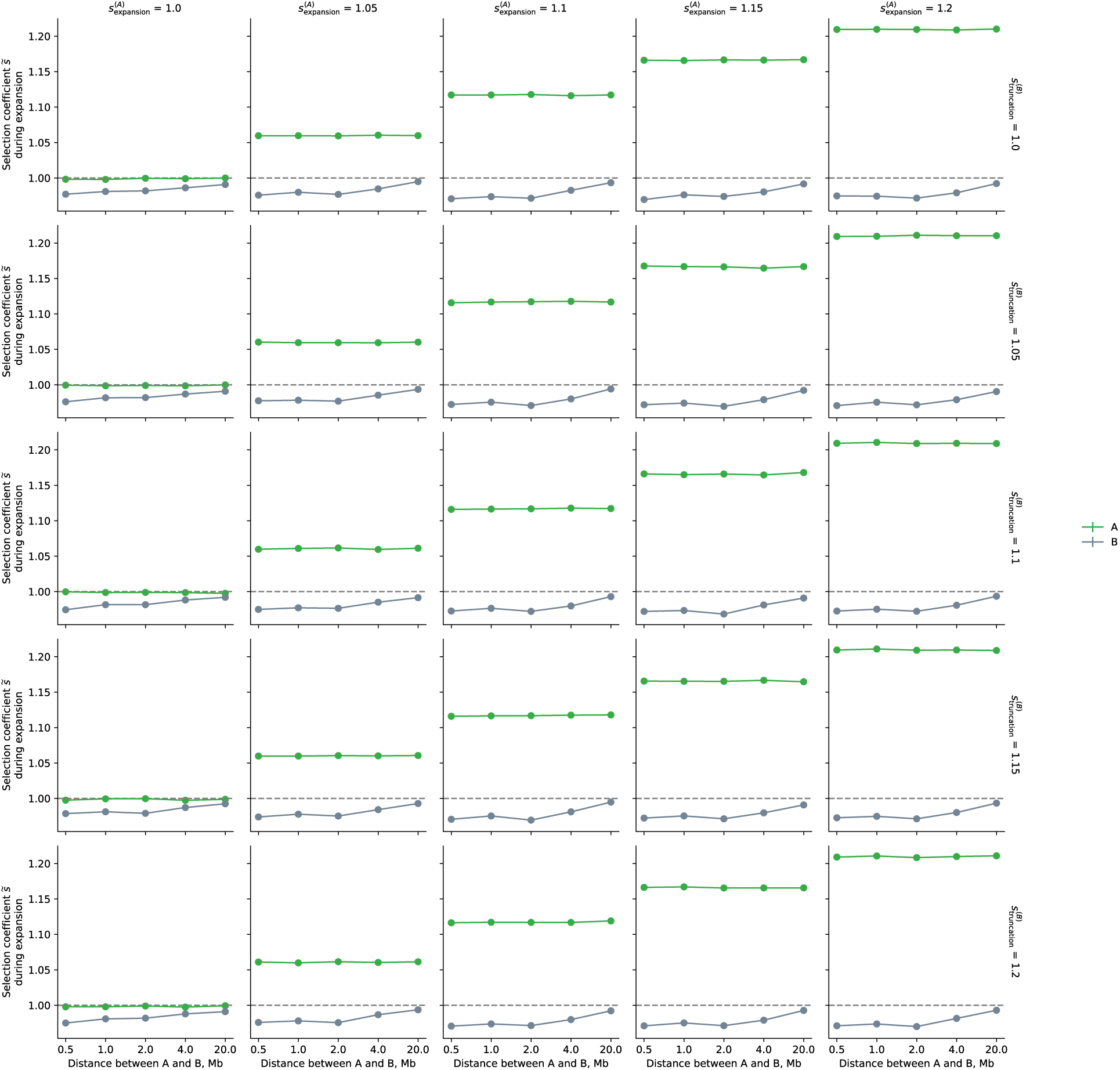
Fitness values (empirical selection coefficients) for two, unlinked alleles during the first generation of truncation, plotted as a function of distance between the two SNPs. Points denote the bootstrapped mean across simulations (N = 250 total simulations) with vertical lines the 95% confidence intervals (CI’s often too small to be visible in plot). The ‘A’ allele was under directional selection during the expansion period (generations 0-9) and neutral during the truncation phase (generations 10-13). The ‘B’ allele was neutral during expansion, then comes under directional selection during truncation. The selection coefficient of ‘A’ was ranged between 1.0 and 1.2 during expansion (columns) and remains 1.0 during truncation. The selection coefficient of ‘B’ was constant at 1.0 during expansion, and ranged between 1.0 and 1.2 during truncation (rows).

**Figure S17.**
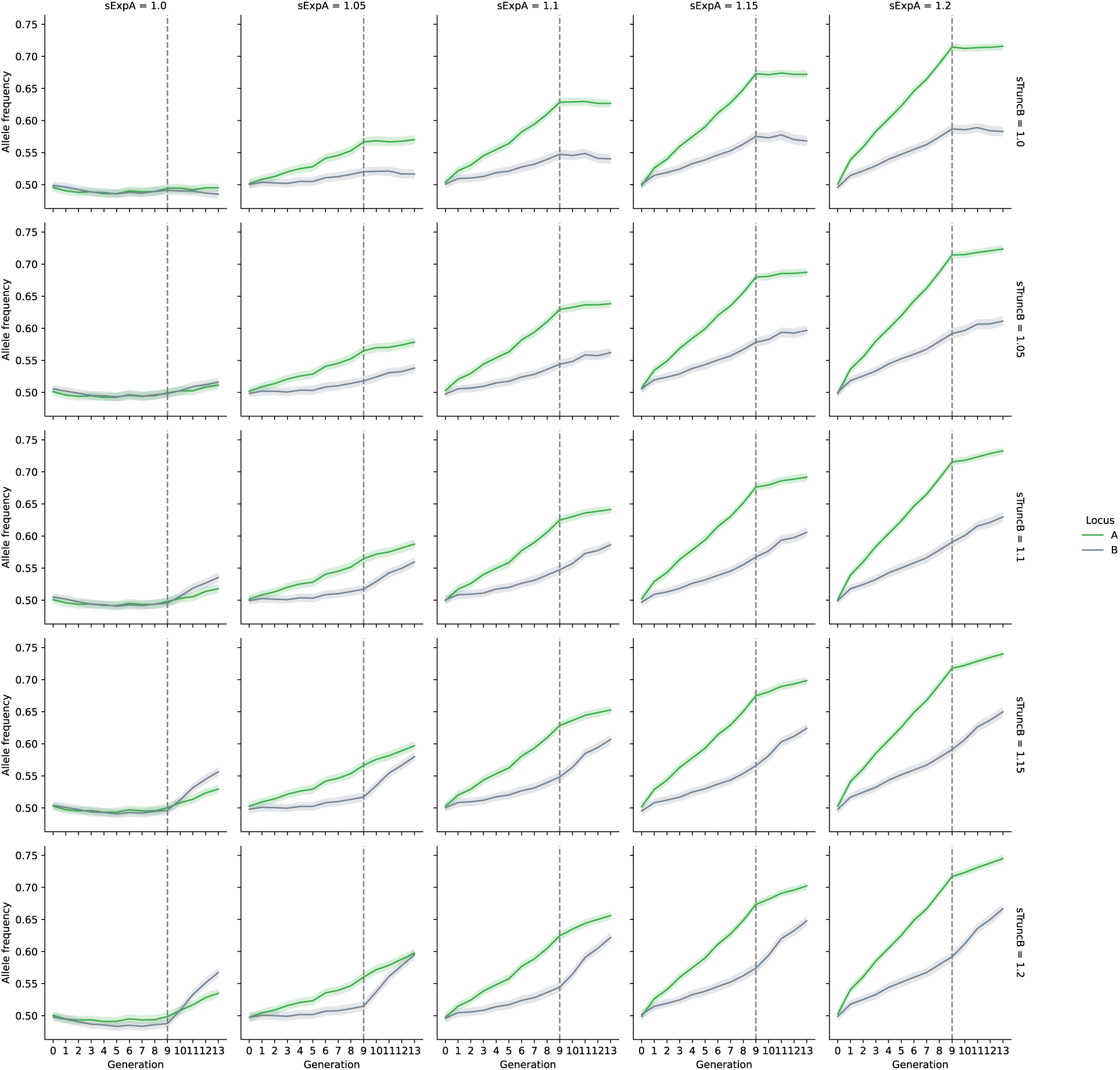
Simulated allele frequency trajectories for two highly linked loci in a state of attraction. The ‘A’ allele is encoded to be under directional selection during the expansion period (generations 0-9) and neutral during the truncation phase (generations 10-13). The ‘B’ allele is encoded neutral during expansion, then comes under directional selection during truncation. The selection coefficient of ‘B’ is constant at 1.0 during expansion and ranged between 1.0 and 1.2 during truncation (rows). The solid line corresponds to the mean frequency value of each allele across simulations and shaded area to the 95% confidence intervals (N = 250 total simulations per set of parameter values).

**Figure S18.**
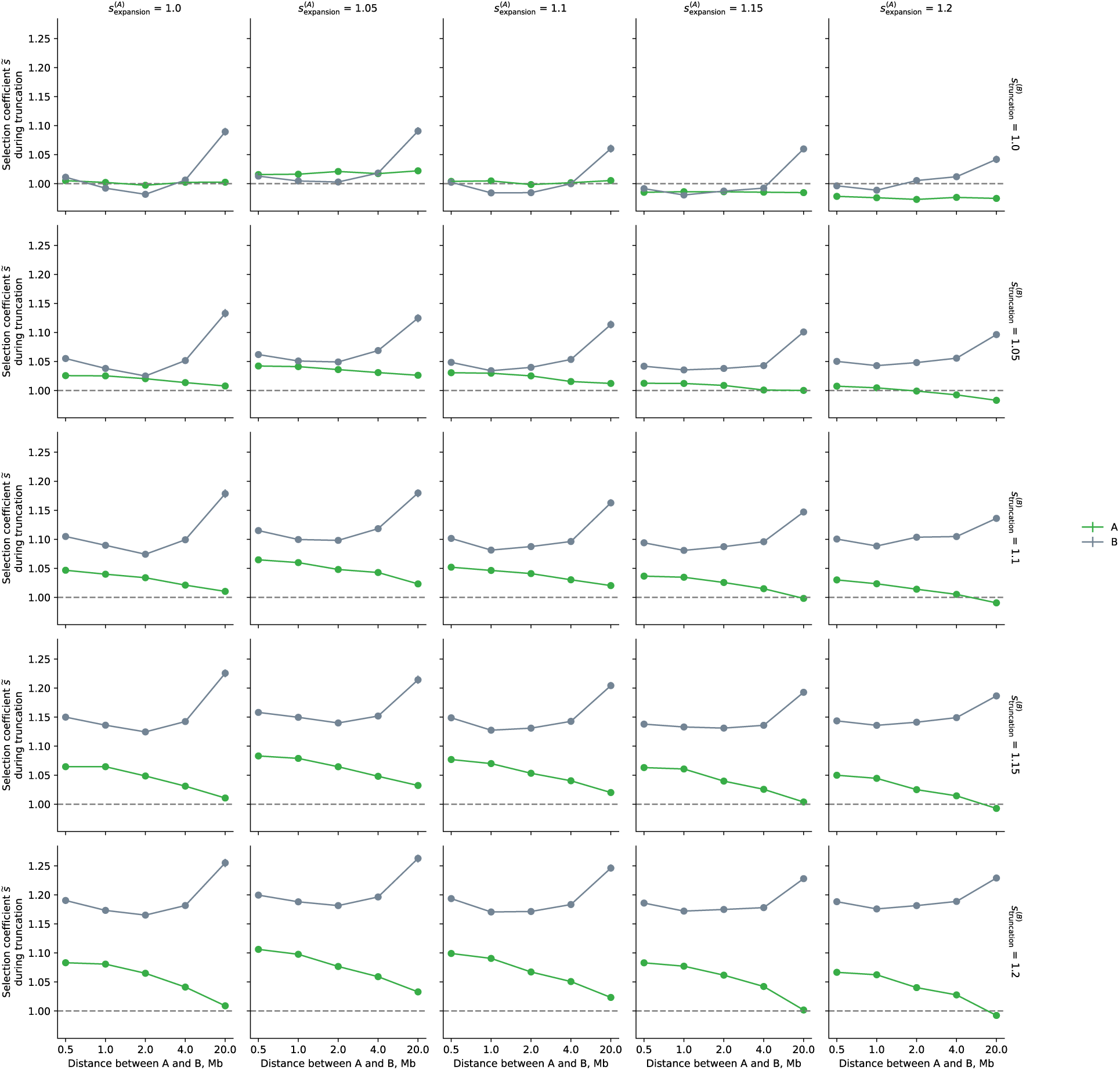
Fitness values (empirical selection coefficients) for two highly linked alleles in a state of attraction, computed during the first generation of truncation and plotted as a function of distance between the two SNPs. Points denote the bootstrapped mean across simulations (N = 250 total simulations) with vertical lines (often not visible in plot) the 95% confidence intervals. The ‘A’ allele was under directional selection during the expansion period (generations 0-9) and neutral during the truncation phase (generations 10-13). The ‘B’ allele was neutral during expansion, then comes under directional selection during truncation. The selection coefficient of ‘A’ was ranged between 1.0 and 1.2 during expansion (columns) and remains 1.0 during truncation. The selection coefficient of ‘B’ was constant at 1.0 during expansion, and ranged between 1.0 and 1.2 during truncation (rows).

**Figure S19.**
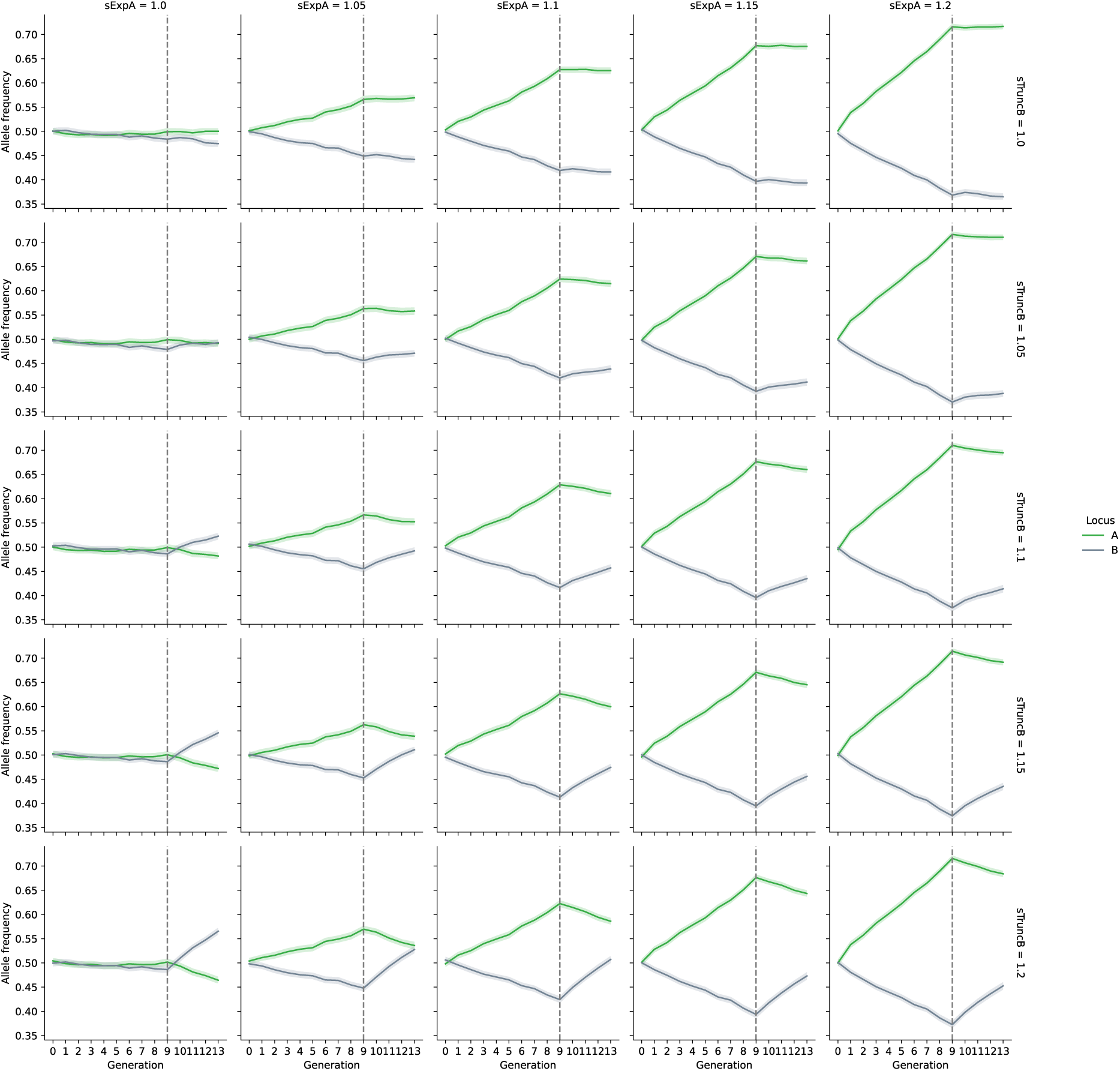
Simulated allele frequency trajectories for two highly linked loci in a state of repulsion. The ‘A’ allele is encoded to be under directional selection during the expansion period (generations 0-9) and neutral during the truncation phase (generations 10-13). The ‘B’ allele is encoded neutral during expansion, then comes under directional selection during truncation. The selection coefficient of ‘B’ is constant at 1.0 during expansion and ranged between 1.0 and 1.2 during truncation. The solid line corresponds to the mean frequency value of each allele across simulations and shaded area to the 95% confidence intervals (N = 250 total simulations per set of parameter values).

**Figure S20.**
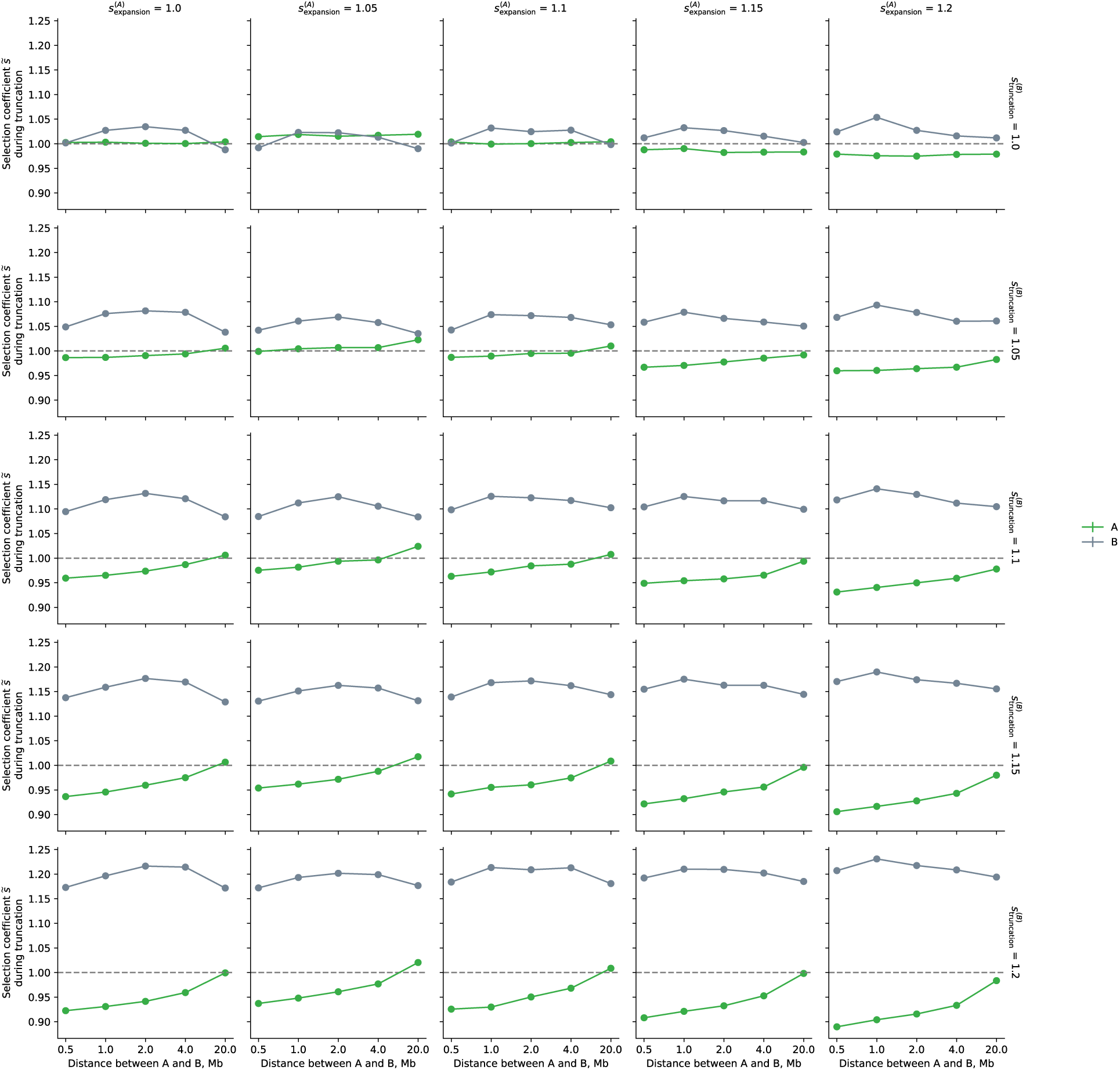
Fitness values (empirical selection coefficients) for two highly linked alleles in a state of repulsion, computed during the first generation of truncation and plotted as a function of distance between the two SNPs. Points denote the bootstrapped mean across simulations (N = 250 total simulations) with vertical lines the 95% confidence intervals (CI’s often not visible in plot). The ‘A’ allele was under directional selection during the expansion period (generations 0-9) and neutral during the truncation phase (generations 10-13). The ‘B’ allele was neutral during expansion, then comes under directional selection during truncation. The selection coefficient of ‘A’ was ranged between 1.0 and 1.2 during expansion (columns) and remains 1.0 during truncation. The selection coefficient of ‘B’ was constant at 1.0 during expansion, and ranged between 1.0 and 1.2 during truncation (rows).

**Figure S21.**
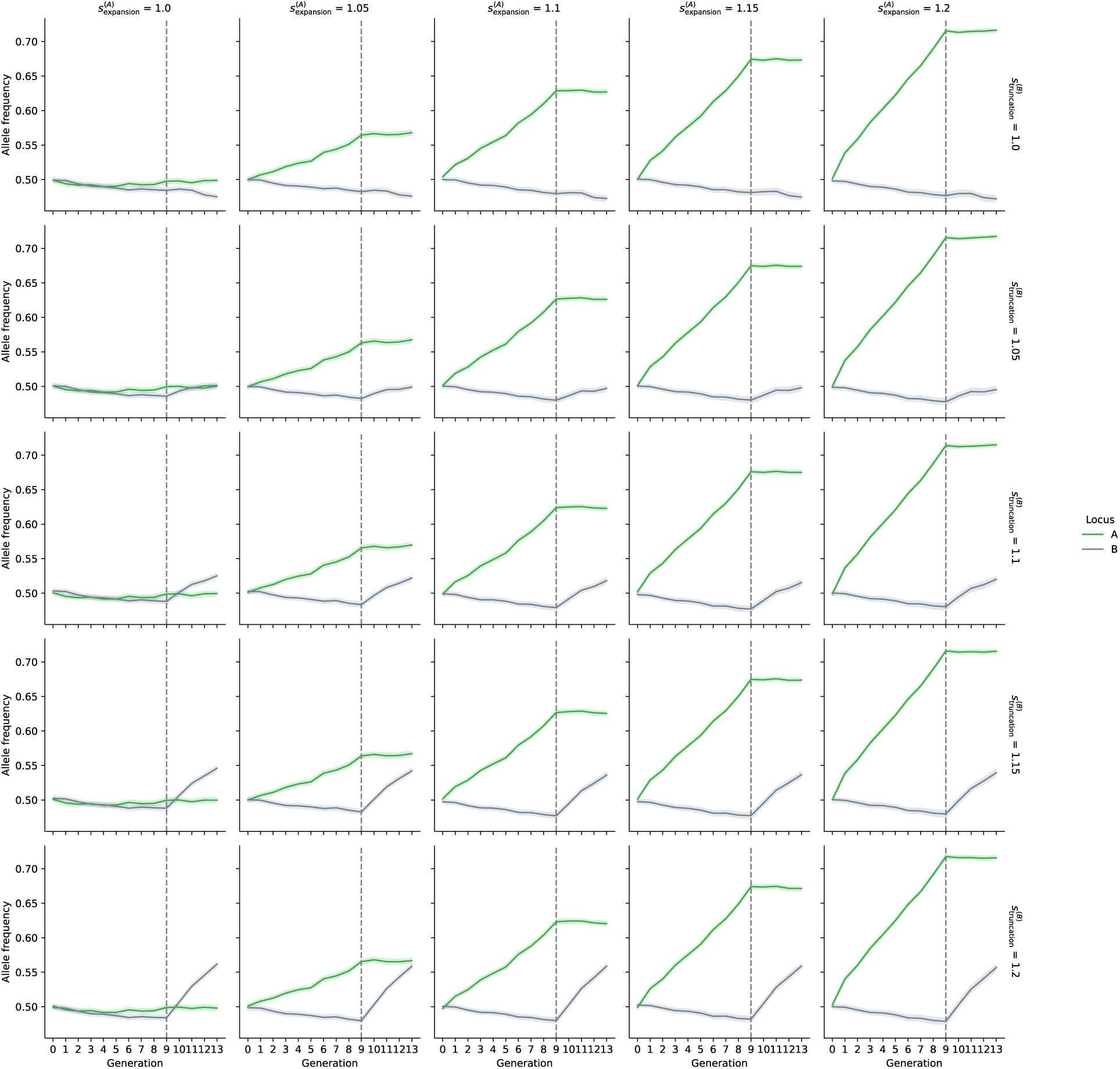
Allele frequency trajectories for two highly linked SNP’s, with selected alleles at each SNP being independently assorted with each other. The ‘A’ allele is encoded to be under directional selection during the expansion period (generations 0-9) and neutral during the truncation phase (generations 10-13). The ‘B’ allele is encoded neutral during expansion, then comes under directional selection during truncation. The selection coefficient of ‘B’ is constant at 1.0 during expansion and ranged between 1.0 and 1.2 during truncation. The solid line corresponds to the mean frequency value of each allele across simulations and shaded area to the 95% confidence intervals (N = 250 total simulations per set of parameter values).

**Figure S22.**
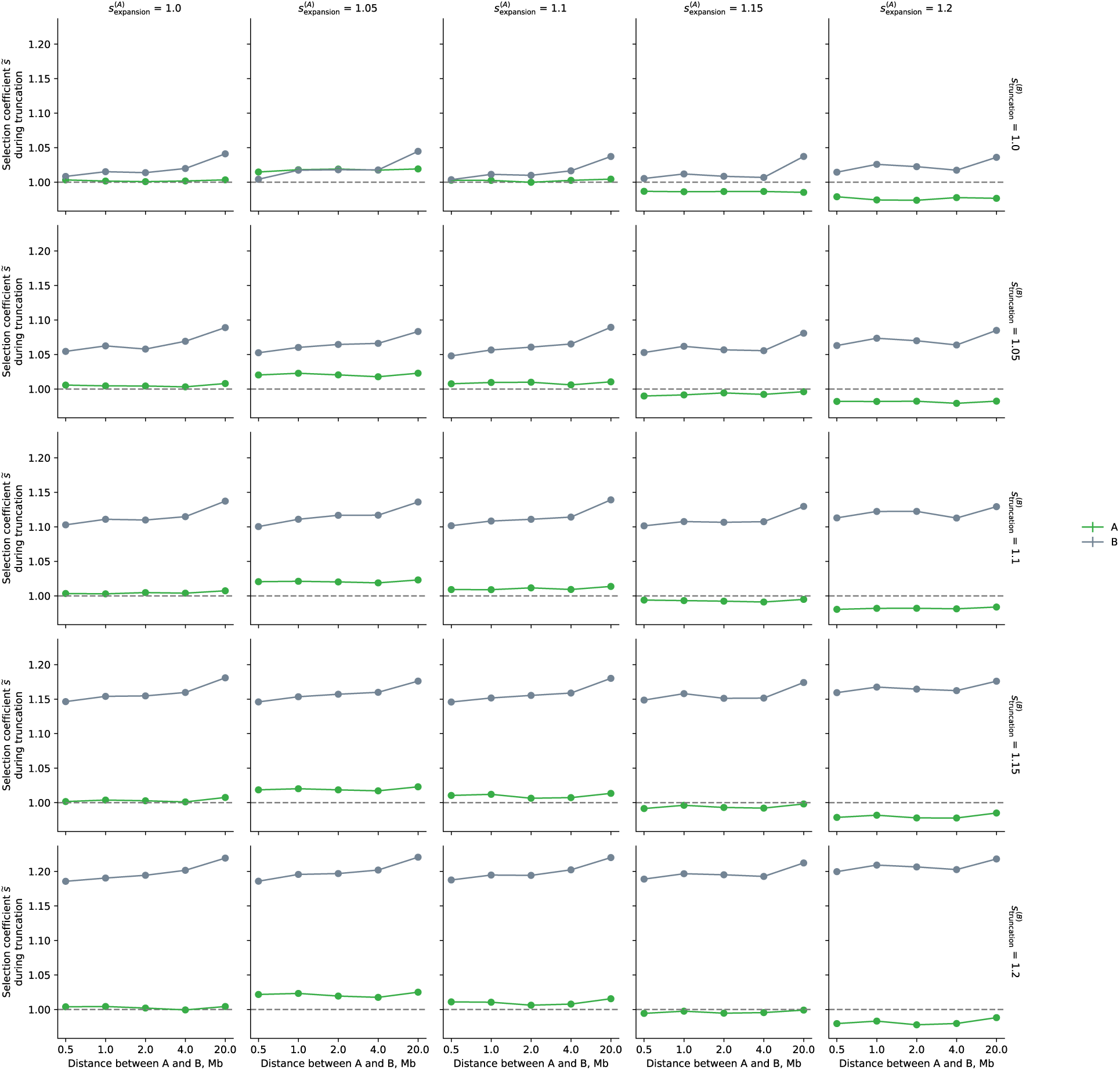
Fitness values (empirical selection coefficients) during the first generation of truncation for the scenario in which the advantageous alleles at linked SNPs (‘A’ and ‘B’) are independently assorted with each other. Points denote the bootstrapped mean across simulations (N = 250 total simulations) with vertical lines (often not visible in plot) the 95% confidence intervals. The ‘A’ allele was under directional selection during the expansion period (generations 0-9) and neutral during the truncation phase (generations 10-13). The ‘B’ allele was neutral during expansion, then comes under directional selection during truncation. The selection coefficient of ‘A’ was ranged between 1.0 and 1.2 during expansion (columns) and remains 1.0 during truncation. The selection coefficient of ‘B’ was constant at 1.0 during expansion, and ranged between 1.0 and 1.2 during truncation (rows).

**Figure S23.**
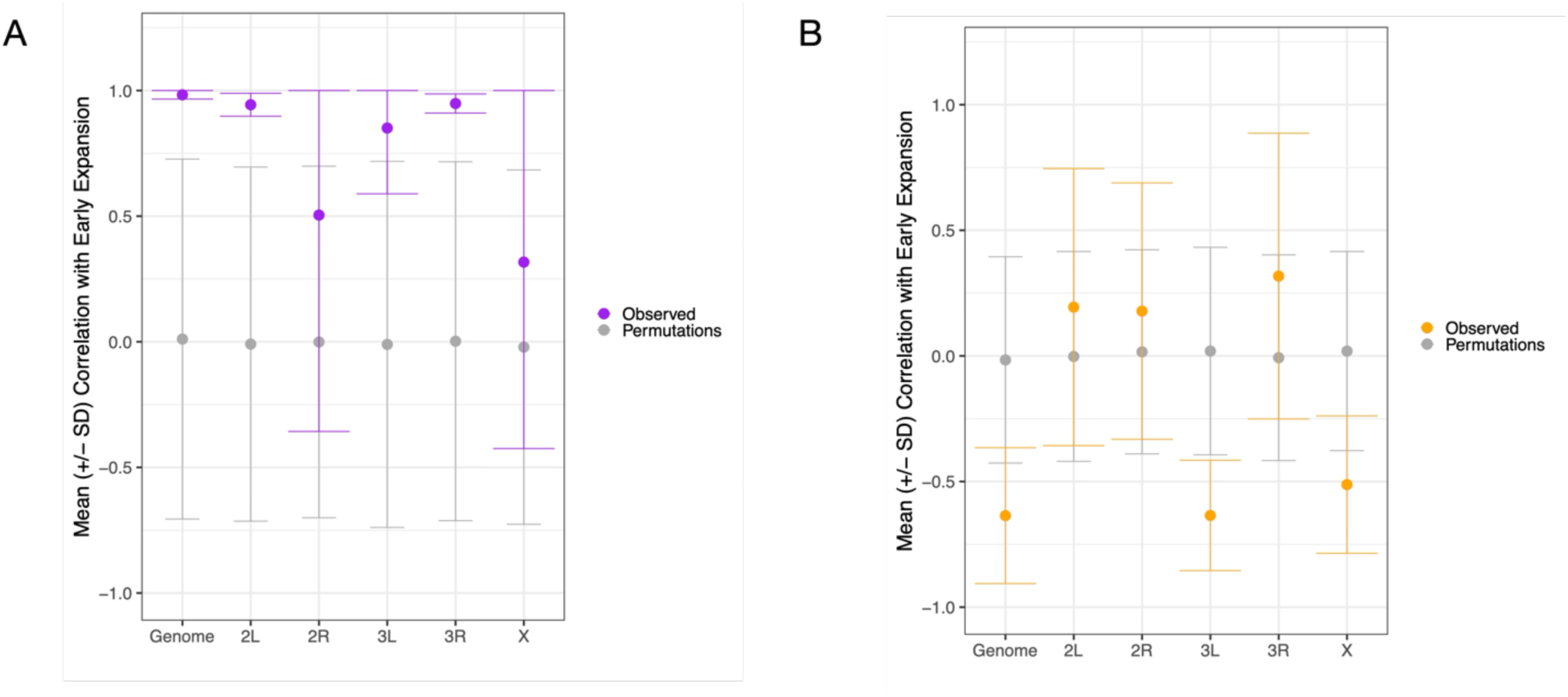
Average Pearson Correlation across replicates (+/- standard deviation) between sample expansion generation and distance along a one-dimensional axis constructed using F_ST_ MDS coordinates for early expansion (generation 0-4) samples. Here, Fst values were derived from SNPs excluding those within, or up to 100 Kb away from, inversion breakpoints. Purple points and error bars correspond to empirical values, while grey points and error bars correspond to values derived from empirical permutations (N = 100).

**Figure S24.**
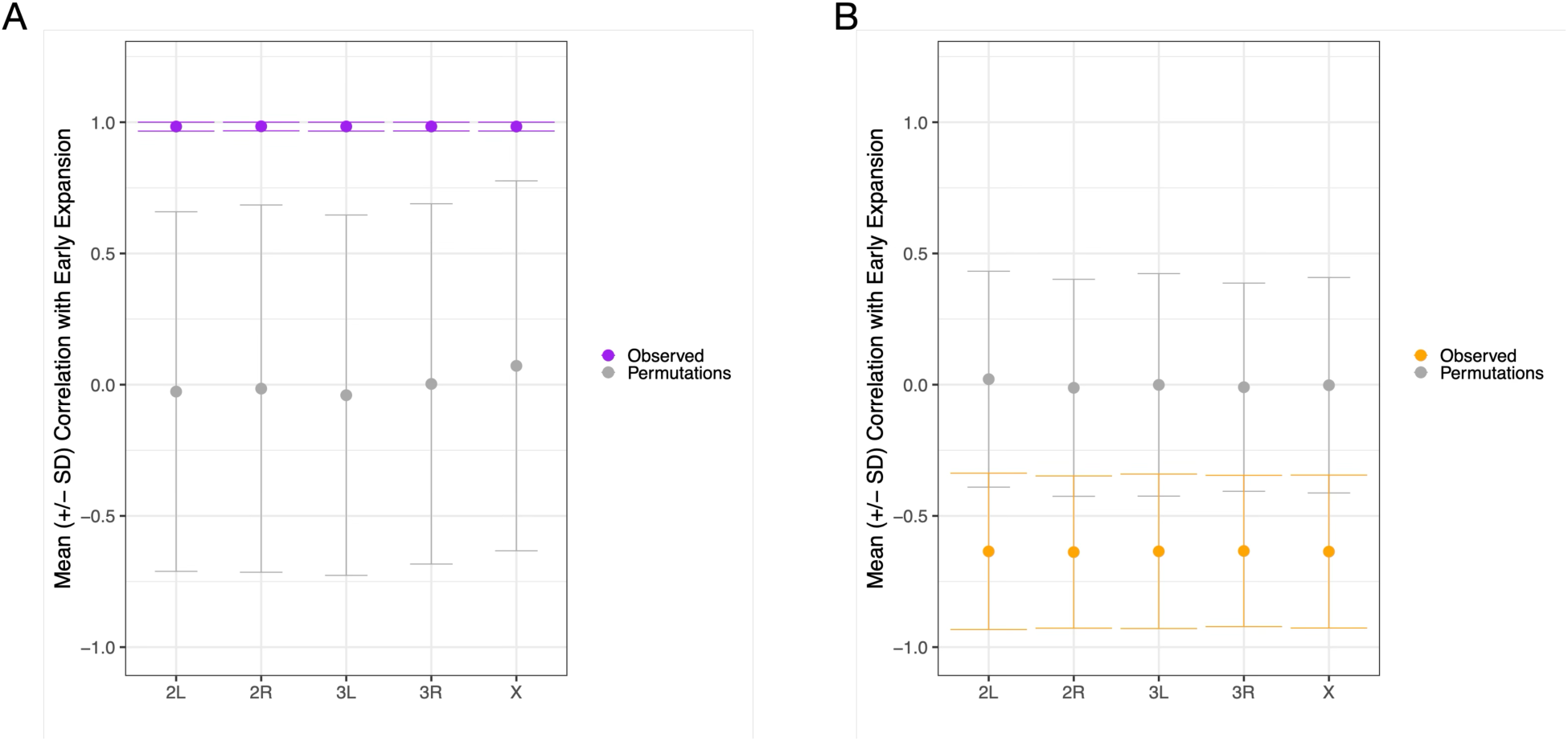
Fst-based MDS analysis derived from randomly sampled SNPs throughout the genome matched to the total number of SNPs present on each chromosomal arm and excluding those within, or up to 100 Kb away from, inversion breakpoints. Plotted are average Pearson Correlation across replicates (+/- standard deviation) between sample expansion generation and distance along a one-dimensional axis constructed using F_ST_ MDS coordinates for early expansion (generation 0-4) samples. Purple points and error bars correspond to empirical values, while grey points and error bars correspond to values derived from empirical permutations (N = 100).

**Figure S25.**
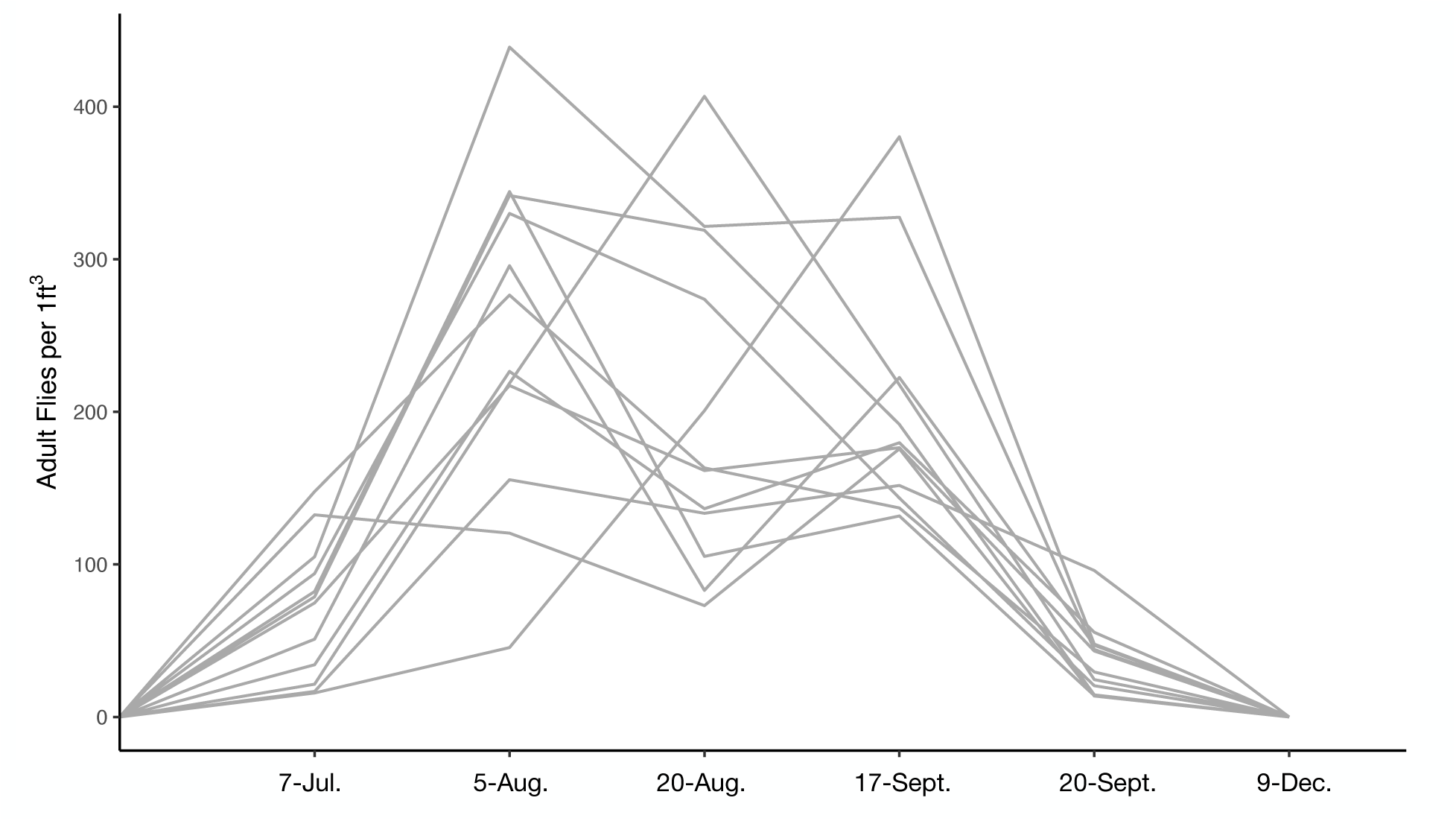
Census estimates of flies in the outdoor mesocosm experiment from 2021. Figure modified from (4).

### Supplementary Tables

**Table S1.**
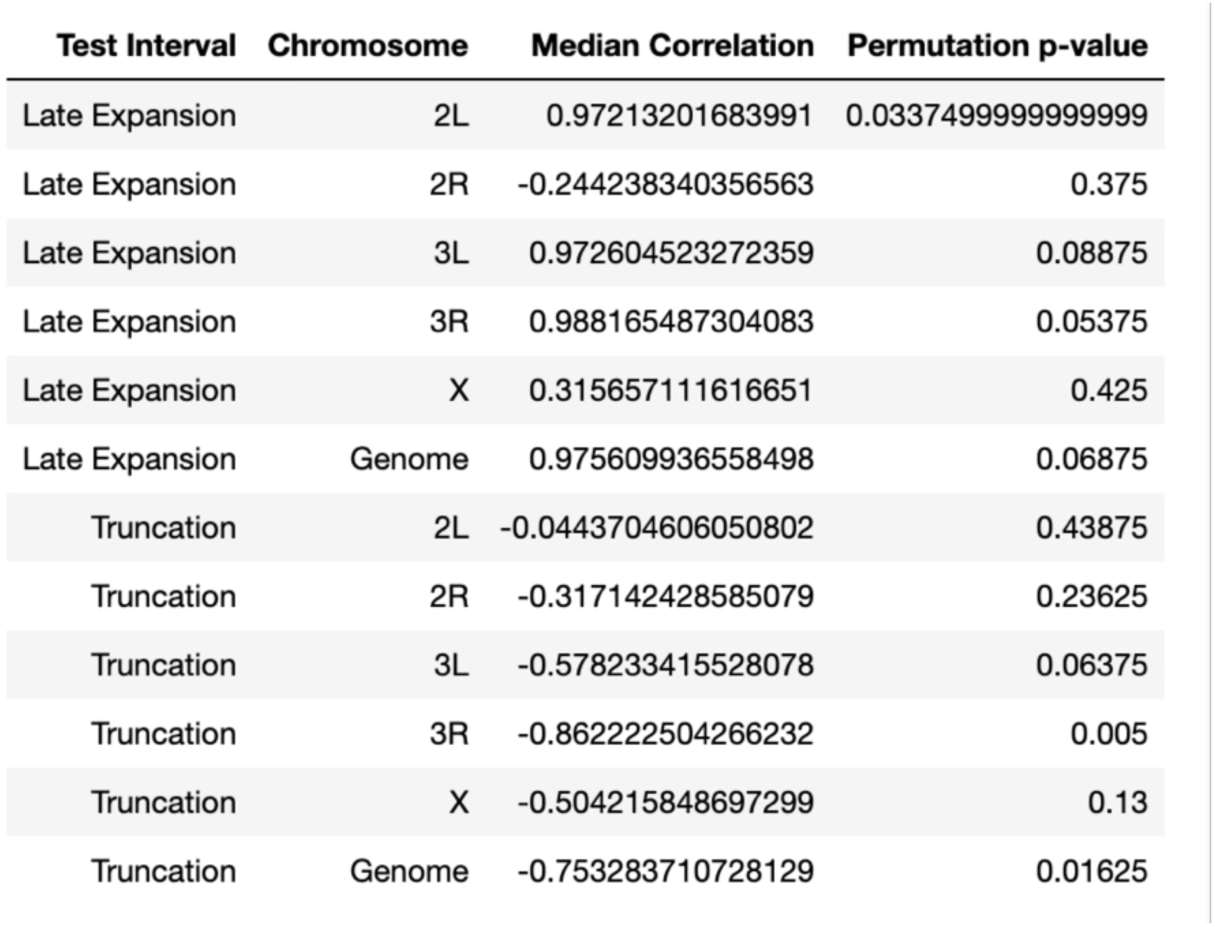
Statistics for projection of late expansion (generations 6-9) and truncation samples onto a one-dimensional axis constructed using F_ST_ MDS coordinates of early expansion (generation 0-4) samples. Correlation coefficients were computed between sample collection time point and distance along this axis for each cage, and reported are the median values across cage for each chromosomal arm, as well as when the analysis was run genome-wide. P-values were generated via comparison of each median correlation to a distribution generated via permutation of time point labels (N = 100 total permutations).

**Table S2.** Statistics for projection of late expansion (generations 6-9) and truncation samples onto a one-dimensional axis constructed using F_ST_ MDS coordinates of early expansion (generation 0-4) samples. Here, F_st_ values and MDS analysis was run on SNPs randomly sampled throughout the genome, matching the number of SNPs present on each chromosomal arm. Correlation coefficients were computed between sample collection time point and distance along this axis for each cage, and reported are the median values across cage for each chromosomal arm, as well as when the analysis was run genome-wide. P-values were generated via comparison of each median correlation to a distribution generated via permutation of time point labels (N = 100 total permutations).

| Test Interval | Chromosome | Median Correlation | Permutation p-value |
| --- | --- | --- | --- |
| Late Expansion | 2L | 0.975716766359953 | 0.07375 |
| Late Expansion | 2R | 0.974541408118333 | 0.075 |
| Late Expansion | 3L | 0.975540423627953 | 0.01875 |
| Late Expansion | 3R | 0.9752928069539 | 0.02625 |
| Late Expansion | X | 0.973467774354005 | 0.0600000000000001 |
| Truncation | 2L | -0.751876958283634 | 0.02 |
| Truncation | 2R | -0.753904647531668 | 0.01875 |
| Truncation | 3L | -0.7502678134965 | 0.04125 |
| Truncation | 3R | -0.750207616089703 | 0.025 |
| Truncation | X | -0.756320833115252 | 0.0225 |

**Table S3.**
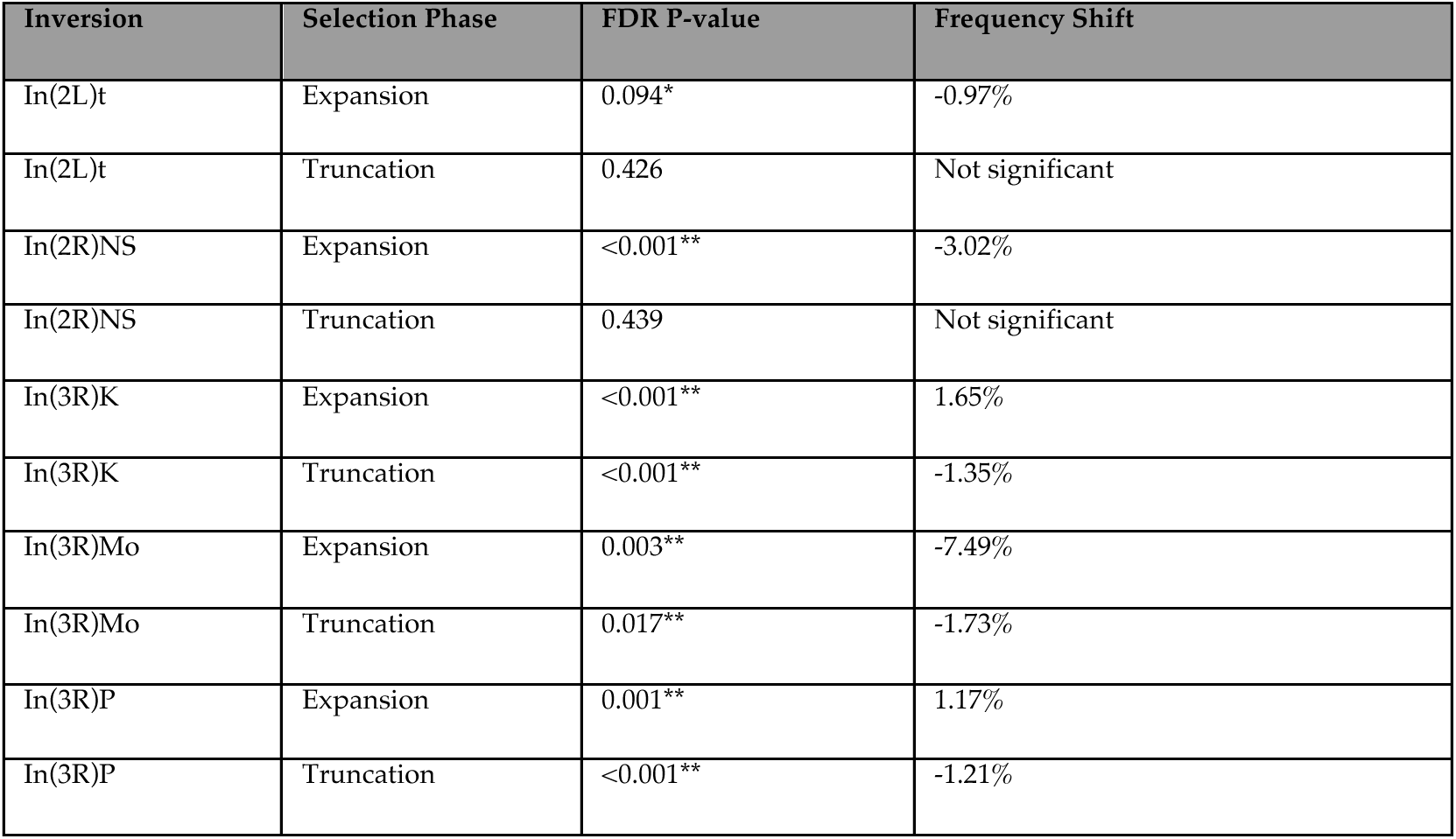
Inversion Frequency Dynamics. GLM regression summary statistics of the expansion and truncation dynamics of four major cosmopolitan inversions segregating in our outbred population of DGRP lines. The FDR corrected p-value for each inversion during both phases of the experiment (expansion and truncation) are presented, alongside the mean inversion frequency shift in cases in which the GLM regression was significant.

**Table S4.**
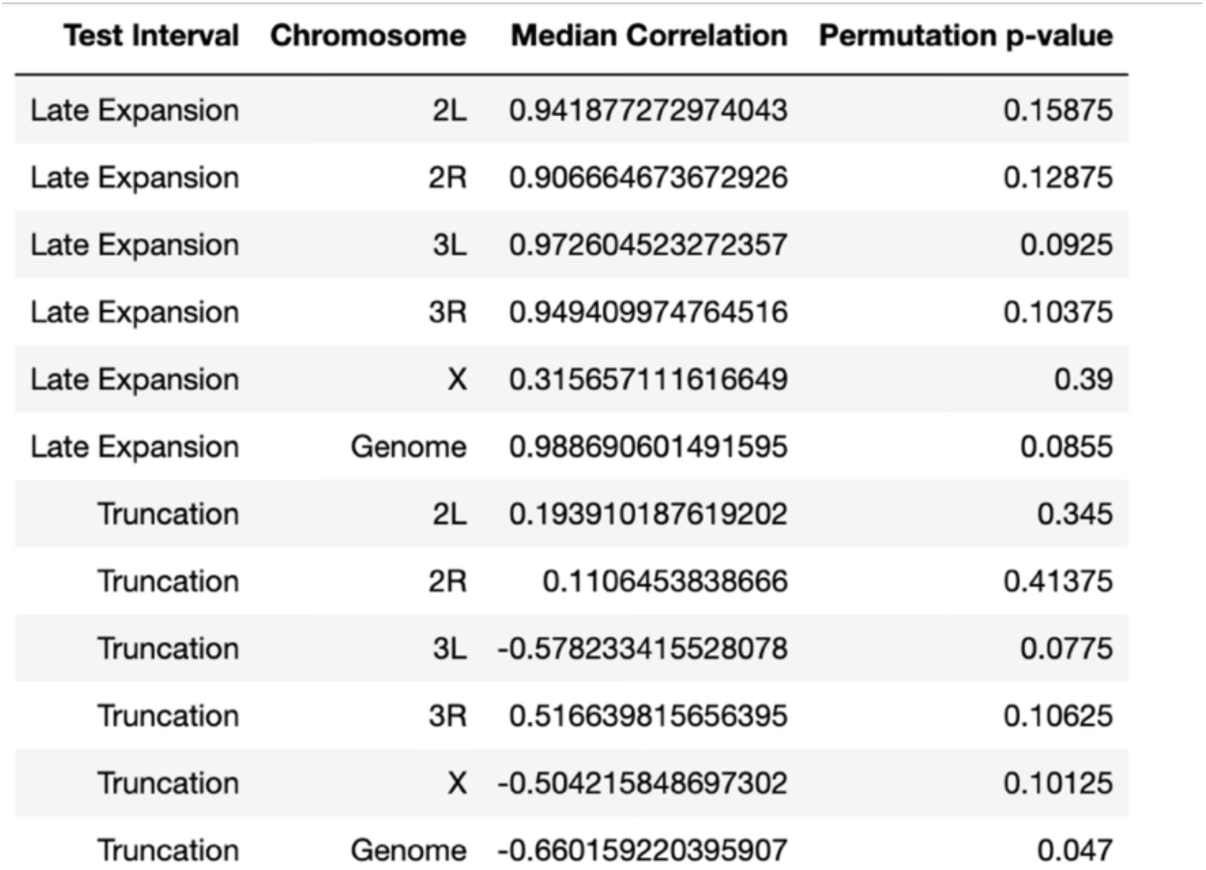
Statistics for projection of late expansion (generations 6-9) and truncation samples onto a one-dimensional axis constructed using F_ST_ MDS coordinates of early expansion (generation 0-4) samples. Here, Fst values and MDS analysis was run on a set of SNPs excluding those within, or up to 100 Kb away from, inversion breakpoints. Correlation coefficients were computed between sample collection time point and distance along this axis for each cage, and reported are the median values across cage for each chromosomal arm, as well as when the analysis was run genome-wide. P-values were generated via comparison of each median correlation to a distribution generated via permutation of time point labels (N = 100 total permutations).

**Table S5.** Statistics for projection of late expansion (generations 6-9) and truncation samples onto a one-dimensional axis constructed using F_ST_ MDS coordinates of early expansion (generation 0-4) samples. Here, Fst values and MDS analysis was run on SNPs randomly sampled throughout the genome, matched to the number of SNPs present on each chromosomal arm, and excluding those within, or up to 100 Kb away from, inversion breakpoints. Correlation coefficients were computed between sample collection time point and distance along this axis for each cage, and reported are the median values across cage for each chromosomal arm, as well as when the analysis was run genome-wide. P-values were generated via comparison of each median correlation to a distribution generated via permutation of time point labels (N = 100 total permutations).

| Test Interval | Chromosome | Median Correlation | Permutation p-value |
| --- | --- | --- | --- |
| Late Expansion | 2L | 0.988413197497469 | 0.0525 |
| Late Expansion | 2R | 0.989942495115707 | 0.05125 |
| Late Expansion | 3L | 0.989390312932461 | 0.05 |
| Late Expansion | 3R | 0.990068206319797 | 0.06375 |
| Late Expansion | X | 0.989855205403952 | 0.08125 |
| Truncation | 2L | -0.665094200789921 | 0.04375 |
| Truncation | 2R | -0.661142450550023 | 0.055 |
| Truncation | 3L | -0.660888966235905 | 0.0525 |
| Truncation | 3R | -0.653623292619365 | 0.03875 |
| Truncation | X | -0.660601423436917 | 0.045 |

**Table S6.** Statistics for comparison of rising allele frequency shifts of indoor-identified SNPs in an outdoor mesocosm during population expansion and collapse. Target values (Mean.Target, Median.Target) indicate those derived from indoor-identified SNPs, while matched values indicate those derived from control SNPs. The distributions of target and matched sites were compared using both a two-tailed t-test and sign test. The resulting test statistics are provided.

| Chromosome | Mean.Target | Median.Target | Mean.Matched | Median.Matched | t | FDR.t.test | FDR.sign.test | Test.Interval |
| --- | --- | --- | --- | --- | --- | --- | --- | --- |
| 2L | 0.0319985981378007 | 3.038013e-02 | 0.00316831802589668 | 3.178635e-03 | 90.4650925090862 | 0.000000e+00 | 8.326673e-16 | Expansion |
| 2R | 0.028969065444205 | 2.551098e-02 | 0.00115295987609255 | 1.615365e-03 | 66.0420368142833 | 0.000000e+00 | 8.326673e-16 | Expansion |
| 3L | 0.0335003699797639 | 3.080976e-02 | 0.00286597441953921 | 3.928358e-03 | 90.8841932663739 | 0.000000e+00 | 8.326673e-16 | Expansion |
| 3R | 0.0369291946950844 | 3.378144e-02 | 0.00264119051901511 | 3.484494e-03 | 129.54052005962 | 0.000000e+00 | 8.326673e-16 | Expansion |
| X | 0.0288667950506608 | 2.644817e-02 | 0.0025943771973813 | 3.045992e-03 | 60.462689719572 | 0.000000e+00 | 8.326673e-16 | Expansion |
| Genome | 0.0333054320018882 | 3.057965e-02 | 0.00260823894732216 | 3.234185e-03 | 199.627700493635 | 0.000000e+00 | 8.326673e-16 | Expansion |
| 2L | -0.00487214737932314 | -4.692767e-03 | -0.00104825093438945 | -9.152650e-04 | -19.5690789120401 | 2.356927e-84 | 7.180716e-52 | Collapse |
| 2R | 0.000478905306333812 | 8.868033e-04 | -0.000171925992292266 | 4.171333e-05 | 2.8948882077324 | 3.958655e-03 | 9.993639e-05 | Collapse |
| 3L | 0.00187994456347232 | 1.696414e-03 | -7.9971808333333e-05 | 2.173192e-04 | 8.39801534836484 | 5.830216e-17 | 1.057355e-15 | Collapse |
| 3R | -0.000895085879399349 | -1.074300e-04 | 0.000631816786767623 | 6.350982e-04 | -8.12116003302136 | 5.475990e-16 | 3.364728e-06 | Collapse |
| X | -0.000821591180249633 | 4.711083e-05 | -0.000622166078071464 | 3.165075e-04 | -0.573004050626995 | 5.666659e-01 | 3.673839e-02 | Collapse |
| Genome | -0.00109679281216485 | -5.765000e-04 | -0.000135973486759733 | 9.451667e-05 | -9.42612164714719 | 4.364745e-21 | 2.861040e-08 | Collapse |

**Table S7.**
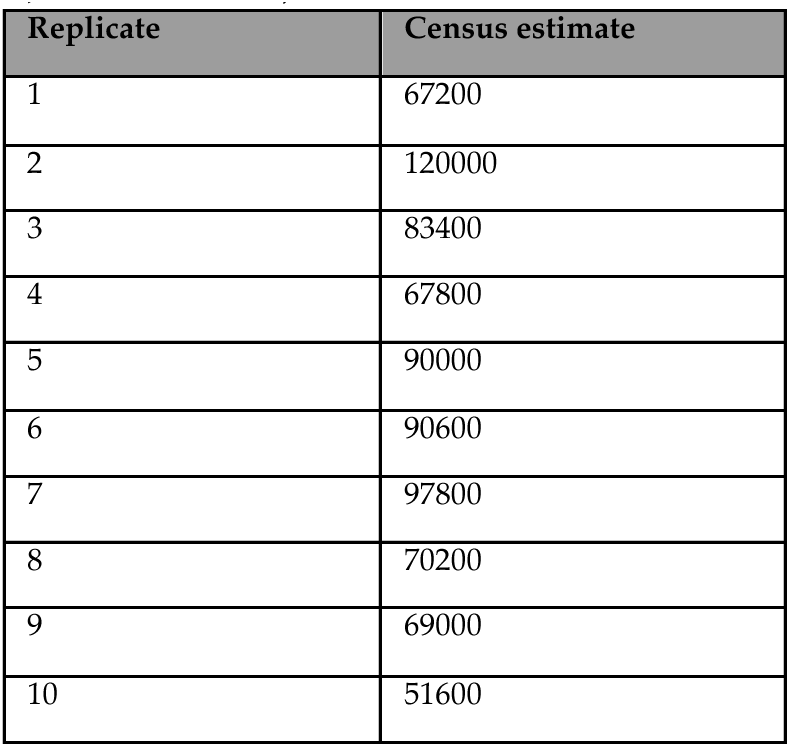
Final fly census size in indoor cages from paired indoor-outdoor mesocosm study. Census sizes were estimated volumetrically using the total remaining flies at the final collection time point (each replicate was originally seeded with 1,000 total flies).

## Notes

### Competing Interest Statement

The authors have declared no competing interest.

### Summary of Updates

The key revision associated with this version of the mansucript is an extensive simulation-based analysis to infer the extent to which the observed patterns of allele frequency change are attributable to antagonistic pleiotropy at a single SNP and/or antagonistic pleiotropy at the haplotype-level.

